# Matched primary-metastasis single-nucleus and spatial profiling of leiomyosarcoma identifies antigen-presenting fibroblasts at the tumor-organ interface

**DOI:** 10.64898/2026.10.02.754332

**Authors:** Anna Kübler, Linda Bergmayr, Thomas Knösel, Philipp Jurmeister, Yauheniya Zhdanovich, Luc M. Berclaz, Josephine Deleuran Hendriksen, Dilara Karaoglu, Romy L. Zuber, Michael Hoberger, Dorit Di Gioia, Markus Albertsmeier, Matthias Grube, Martin Trepel, Carolin Mogler, Friedrich Anger, Katja Fechner, Philipp Keyl, Stefan Fröhling, Frederick Klauschen, Lars H. Lindner, Anton Burkhard-Meier, Andreas Mock

## Abstract

Leiomyosarcoma is an aggressive soft tissue sarcoma that metastasizes preferentially to lung and liver. Because matched primary and metastatic specimens are rarely available, the cellular changes that accompany metastatic progression remain poorly defined. Here we combined single-nucleus RNA sequencing of archival FFPE tissue from four matched primary-metastasis pairs (two lung, two liver metastases; 48,941 nuclei) with imaging-based spatial transcriptomics of an extended tissue microarray cohort of 21 patients (69 cores, 313,218 cells). Malignant cells, not assignable from transcriptomic references alone, were defined by inferred copy number profiles that were concordant with matched DNA methylation arrays and confirmed the clonal origin of each metastasis. Malignant metaprograms were organ-associated, with a stress- and hypoxia-associated program enriched in lung metastases and a proliferative, matrix-remodeling program enriched in liver metastases. Antigen-presenting cancer-associated fibroblasts (apCAFs) were enriched in metastases relative to matched primaries. This shift was reproduced in the spatial cohort (p = 0.001), where apCAFs formed a discrete band at the tumor-organ interface between malignant cells and adjacent parenchyma, whereas they were sparse and randomly distributed in primary tumors. Spatial transcriptomics further resolved the malignant compartment into three histomolecular phenotypes: (i) smooth-muscle-like with myxoid features, (ii) GPNMB⁺ with anaplastic features, and (iii) PDGFRB⁺ with vessel-rich stroma, each with a concordant immunohistochemical (α-SMA, GPNMB, PDGFRB) and morphological correlate. Together, our data provide a matched primary-metastasis reference for leiomyosarcoma, identify apCAFs as a spatially organized, metastasis-associated stromal population, and delineate histomolecular phenotypes recognizable by routine immunohistochemistry, including a GPNMB⁺ phenotype potentially amenable to GPNMB-directed therapies.

## Introduction

Sarcomas comprise a heterogeneous group of rare malignancies arising from mesenchymal tissue, accounting for approximately 1% of adult cancers^1^. Their rarity and biological diversity pose substantial challenges for both clinical management and research^2^. Sarcomas are broadly classified as bone or soft tissue tumors^3^, among the latter, leiomyosarcoma (LMS) is one of the most prevalent subtypes, constituting approximately 10-20% of soft tissue sarcoma cases^3^. LMS is defined by smooth muscle differentiation and occurs at multiple anatomical sites, including the uterus, retroperitoneum, large blood vessels and extremities. It is an aggressive malignancy characterized by a high risk of recurrence and metastatic spread, most frequently to the lung and liver^4^. More than half of patients develop metastatic disease, and outcomes for advanced LMS remain poor. While surgical resection remains the cornerstone of treatment for localized disease, metastatic LMS is primarily treated with anthracycline-based systemic therapy^5^. Oligometastatic disease is considered an intermediate stage between localized disease and widespread metastatic dissemination, defined by a limited metastatic burden that may still be amenable to local treatment with curative intent^6,7^. In selected patients with oligometastatic soft tissue sarcoma, long-term survival may be achieved through metastasis-directed local treatments^8^. Prognostic and predictive factors that could guide patient selection for such treatment are, however, lacking, and the biology that distinguishes an LMS metastasis from its primary tumor is largely unknown.

At the level of malignant cells, LMS is characterized by complex karyotypes with recurrent losses in TP53, RB1 and PTEN rather than by recurrent driver fusions^9^. Primary and metastatic tumors may differ fundamentally in their growth drivers and in the composition of their tumor microenvironment (TME)^10^. While primary tumors are shaped largely by their tissue of origin, disseminated cells must adapt to the constraints of a foreign organ, including immune surveillance, hypoxia, and altered nutrient availability^11^. Which of these constraints determines whether LMS cells establish in lung or liver - the two dominant metastatic sites - is not understood, and the two niches are rarely examined side by side. Paired cohorts of LMS profiled at single-cell resolution have, to our knowledge, not been reported.

Within the TME, cancer-associated fibroblasts (CAFs) are central mediators of tumor-stroma crosstalk, acting through secreted factors, direct cell-cell contact, extracellular matrix (ECM) remodeling, and immune modulation^12,13^. CAFs are highly heterogeneous and plastic, and distinct subtypes exert tumor-promoting or tumor-restraining effects depending on tumor type, disease stage, and microenvironmental context^14,15^. Single-cell studies, conducted predominantly in carcinomas, have resolved multiple CAF subtypes, including myofibroblastic (myCAFs), inflammatory (iCAFs), and antigen-presenting CAFs (apCAFs), the latter defined by expression of MHC class II molecules and CD74 in the absence of classical co-stimulatory molecules^13,16,17^. Whether this CAF taxonomy maps onto sarcomas remains unclear and is conceptually challenging, because malignant cells and stromal fibroblasts share a mesenchymal origin, intrinsically blurring their transcriptional boundary^18^.

Advances in single-cell and spatial transcriptomics now allow cellular heterogeneity and tissue organization to be assessed at high resolution^19,20^. Critically for rare entities, optimized single-nucleus RNA sequencing (snRNA-seq) protocols permit transcriptomic profiling of formalin-fixed, paraffin-embedded (FFPE) tissue samples^21^, which for LMS is often the only tissue available for both a primary tumor and its metastasis resected up to years apart. Combining snRNA-seq with imaging-based spatial transcriptomics additionally preserves the tissue context that determines which cells can physically interact^22^.

Here, we investigated how the cellular composition of LMS changes between primary tumors and matched distant metastases, and whether such changes are spatially organized. Using snRNA-seq of four matched primary-metastasis pairs from FFPE tissue, we generated a finely annotated LMS reference that resolves malignant, stromal, immune and organ-resident compartments. Comparison of CAF subtypes across tumor sites identified apCAFs as enriched in metastases. We then used imaging-based spatial transcriptomics in an extended tissue microarray cohort of 21 patients to validate the apCAF enrichment and show that metastatic apCAFs preferentially accumulate at the interface between tumor and host organ. In the same cohort, the malignant compartment resolved into three histomolecular phenotypes - smooth-muscle-like with myxoid features, GPNMB⁺ with anaplastic features, and PDGFRB⁺ with vessel-rich stroma. Together, these data provide a matched primary-metastasis reference for LMS, identify a spatially defined, metastasis-associated stromal population, and delineate malignant phenotypes recognizable by routine immunohistochemistry.

## Results

### Clinical characteristics of the study cohort

The study cohort comprises 21 patients with metastatic leiomyosarcoma (LMS), with a median age of 54 years at first diagnosis. Lung and liver were the predominant sites of first metastasis, occurring in 12 (57.1%) and 8 (38.1%) patients, respectively (**Table S1**). In total, 35 tumor specimens were analyzed, including 13 primary tumors, two primary tumor recurrences, and 20 distant metastases (**Table S2**). According to the ESTRO-EORTC classification, eight distant metastases were classified as metachronous oligorecurrence (40%), five as induced oligopersistence (25.0%), three as synchronous oligometastatic disease (15%), and two each as repeat oligorecurrence and induced oligoprogression (10% each).

The median follow-up from the diagnosis of first metastasis was 72.5 months (95% CI, 49.2–84.1). Median overall survival (OS), measured from the date of first metastasis, was not reached (95% CI, 48.6 months-not reached). Following the first metastasectomy, median follow-up was 68.5 months (95% CI, 43.4-81.1), with a median progression-free survival (PFS) of 10.0 months (95% CI, 6.8-21.1) and median OS was not reached (95% CI, 43.8 months-not reached). Among the metastatic specimens, 11/20 (55%) had been exposed to systemic therapy prior to tissue sampling. In exploratory univariable analyses, none of the assessed clinical factors were significantly associated with PFS or OS (**Table S3**).

### A single-nucleus reference dataset of paired primary and metastatic leiomyosarcoma

The overall study design is shown in **Figure 1a**. To characterize the cellular composition and transcriptional heterogeneity of LMS at single-cell resolution, we performed snRNA-seq on four matched primary-metastasis pairs, comprising four primary tumors and their corresponding metastases (two lung, two liver; **Fig 1b; Fig. S1a**). All samples were FFPE blocks from routine diagnostics, archived between 2015 and 2020. Following quality control, a total of 48,941 nuclei were retained for downstream analyses. On average 6118 nuclei were recovered per sample (range 3,294 - 9506) with an average sequencing depth of 12,569 reads per nucleus (range 5,744 - 27,152) and 77.8% confidently mapped reads (range 68.4%-87%; **Table S4**).

**Figure 1.**
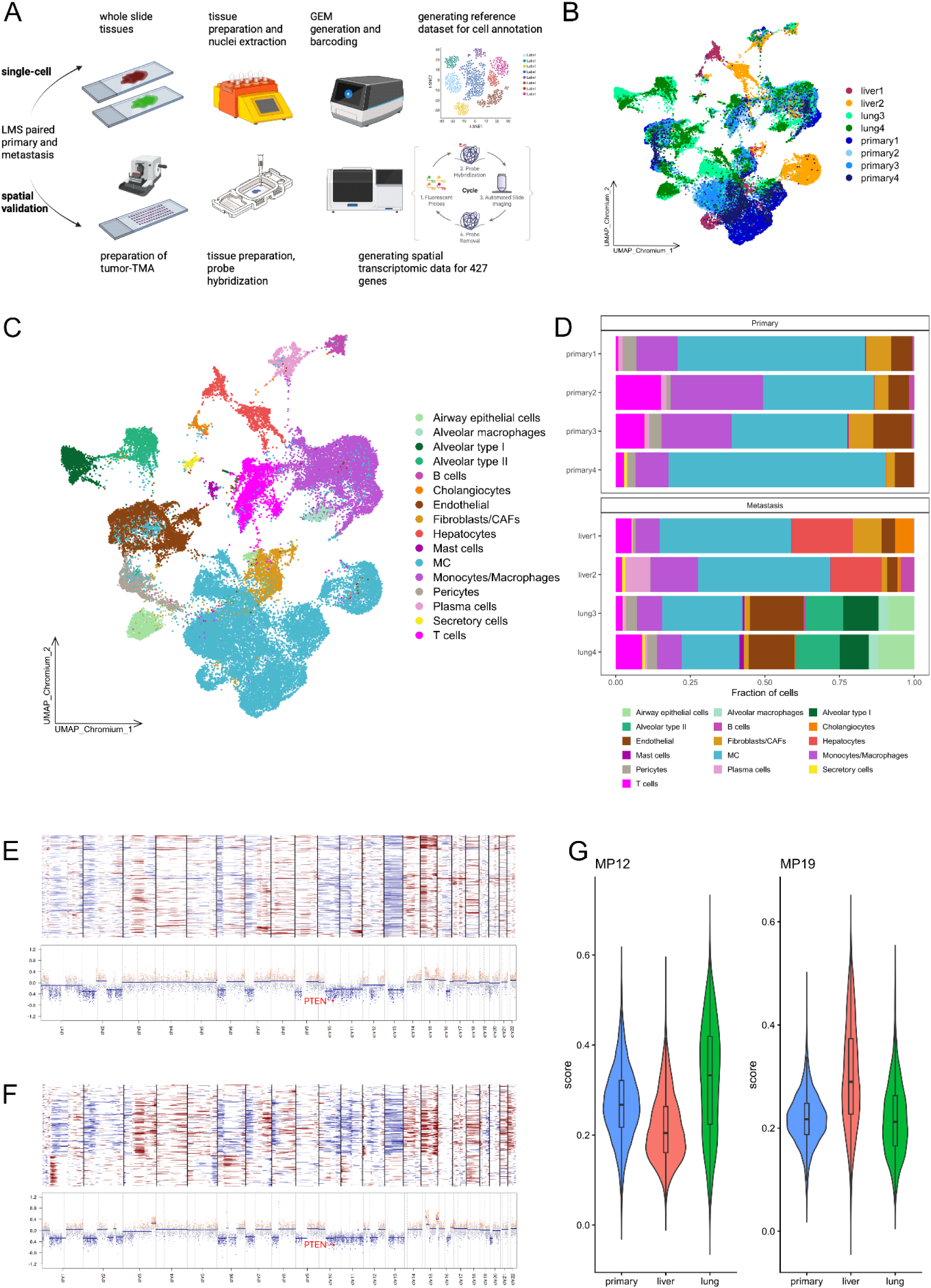
Study design and single-nucleus transcriptomic landscape of paired primary and metastatic LMS. (a) Study design integrating snRNA-seq (Chromium, 10x Genomics) of four matched primary–metastasis pairs with imaging-based spatial transcriptomics (Xenium, 10x Genomics) of a 21-patient tissue microarray. Created in BioRender (Dewenter, I. (2026); https://BioRender.com/akpfadg), incorporating images from the 10x Genomics Support Hub. (b) UMAP embedding of the snRNA-seq data (8 samples), colored by sample. (c) UMAP embedding colored by annotated cell type (n = 48,941 nuclei); MC, malignant cells. (d) Cell type composition per sample, split by primary tumors and metastases. (e, f) Inferred copy number profiles (top) and methylation array-derived copy number profiles (bottom) for a representative primary LMS (e) and its matched metastasis (f); PTEN locus indicated. (g) NMF metaprogram activity scores for MP12 and MP19 across primary tumors, liver metastases and lung metastases.

Louvain-based unsupervised clustering followed by preliminary reference-based annotation identified 16 distinct cell populations, including diverse immune populations (monocytes/macrophages, T cells, B cells, plasma cells, mast cells), stromal populations (fibroblasts/CAFs, endothelial cells, pericytes), lung cells (airway epithelial cells, alveolar type I, alveolar type II, alveolar macrophages), liver cells (hepatocytes, cholangiocytes), secretory cells and one major malignant cell cluster (MC; **Fig. 1c,d**; **Fig. S1b,d**).

Because reference-based annotation alone could not reliably distinguish malignant LMS cells from stromal fibroblasts, which share a mesenchymal transcriptional identity, we inferred copy number variation (CNV) profiles from the single-nucleus expression data, using immune and endothelial cells as non-malignant reference populations. These profiles were validated against DNA methylation array data generated from the same eight samples (**Fig. 1e,f**). This approach enabled discrimination of malignant and non-malignant cell populations (**Fig. 1c**). Chromosomal gains and losses identified by methylation arrays were concordant with the inferred CNV profiles, including, e.g., recurrent loss encompassing the *PTEN* locus. Primary tumors and their matched metastasis displayed highly similar CNV profiles, consistent with a shared clonal origin. Cells with pronounced copy number alterations were consequently classified as malignant, enabling robust discrimination of malignant from non-malignant populations (**Fig. 1c**).

Malignant cells were present in all eight samples and constituted a major population in every sample; immune and stromal cells were likewise detected in every sample (**Fig. 1d**; **Fig. S1b,d**). Overall, malignant cells accounted for an average of 43.3% of all cells, with proportions ranging from 19.3% to 72.7%. The average number of malignant cells per sample was 2542, ranging from 906 to 5015 (**Table S5**). Compared with previously published LMS and sarcoma single-cell datasets (GEO: GSM7634954; GEO: GSE212527), which comprise limited cell numbers and comparatively broad cell type annotations, our dataset provides a higher-resolution cellular reference for LMS. In our data, malignant, stromal, and immune and organ-resident populations could be resolved into distinct subpopulations, and the fully annotated object is provided for reuse (see Data availability).

### Malignant transcriptional programs distinguish lung from liver metastasis

Having defined the malignant compartment, we asked whether malignant cells display site-associated transcriptional states. We applied non-negative matrix factorization (NMF) to the subset of malignant cells and aggregated recurrent gene programs into 20 metaprograms (MPs) (**Fig. S1c**). Five metaprograms (MP1, MP3, MP11, MP14, MP20) were excluded from downstream analysis because activity scores were uniformly low across all samples, leaving 15 MPs that showed substantial activity. The recovered programs corresponded to interpretable biological processes: proliferation and cell cycle (MP4, MP5, MP8, MP19), ECM remodeling (MP6, MP9, MP16, MP17), contractile and smooth muscle identity (MP2, MP10), stress and inflammatory responses (MP7, MP12, MP13, MP15, MP18), and metabolic adaptation (MP10, MP17) (**Fig. S1c**). The persistence of contractile and smooth muscle programs across all samples is consistent with retained lineage identity in LMS.

Comparing metaprogram activity across anatomical sites revealed distinct organ-specific transcriptional adaptations of malignant LMS cells (**Fig. S1e**). Among the identified programs, MP12 showed higher activity in lung metastases than in primary tumors and liver metastases, with a small effect size relative to primary tumors and a large effect size relative to liver metastases (**Fig. 1g**). MP12 was characterized by cellular stress-response genes (NR4A1, KLF6), the hypoxia-associated transcription factor EPAS1, and extracellular matrix-associated genes including TNXB and EMP2. Hallmark gene-set overlap and canonical marker genes therefore annotated MP12 as a stress- and inflammation-associated program with hypoxia adaptation in the pulmonary niche. Conversely, MP19 showed higher activity in liver metastases relative to primary tumors and lung metastases, exhibiting large effect sizes for both comparisons (**Fig. 1g**). MP19 combined proliferation-associated genes (LIG1, HELLS, TYMS, E2F1) with ECM remodeling and contractile genes, including COL6A1, COL6A2, MMP14, ITGA5, MYH11, and MYLK. This profile is consistent with a proliferative, matrix-remodeling malignant cell state associated with hepatic metastases.

Together, these observations indicate that malignant LMS cells retain core smooth muscle features while exhibiting site-associated transcriptional differences in metastatic organs. Specifically, lung metastases were enriched for a stress- and hypoxia-adapted program, whereas liver metastases exhibited a proliferative, matrix-remodeling phenotype.

### CAF subtype composition reveals enrichment of apCAFs in metastases

Fibroblasts and CAFs represented a major stromal population in the LMS microenvironment across all samples. To dissect CAF biology, cells initially annotated as Fibroblasts/CAFs were extracted and re-clustered (**Fig. 2a**). Expression of established CAF subtype marker genes from a published single-cell reference^17^ supported the presence of multiple transcriptionally distinct CAF states in LMS (**Fig. 2d**; **Fig. S2a,d**).

**Figure 2.**
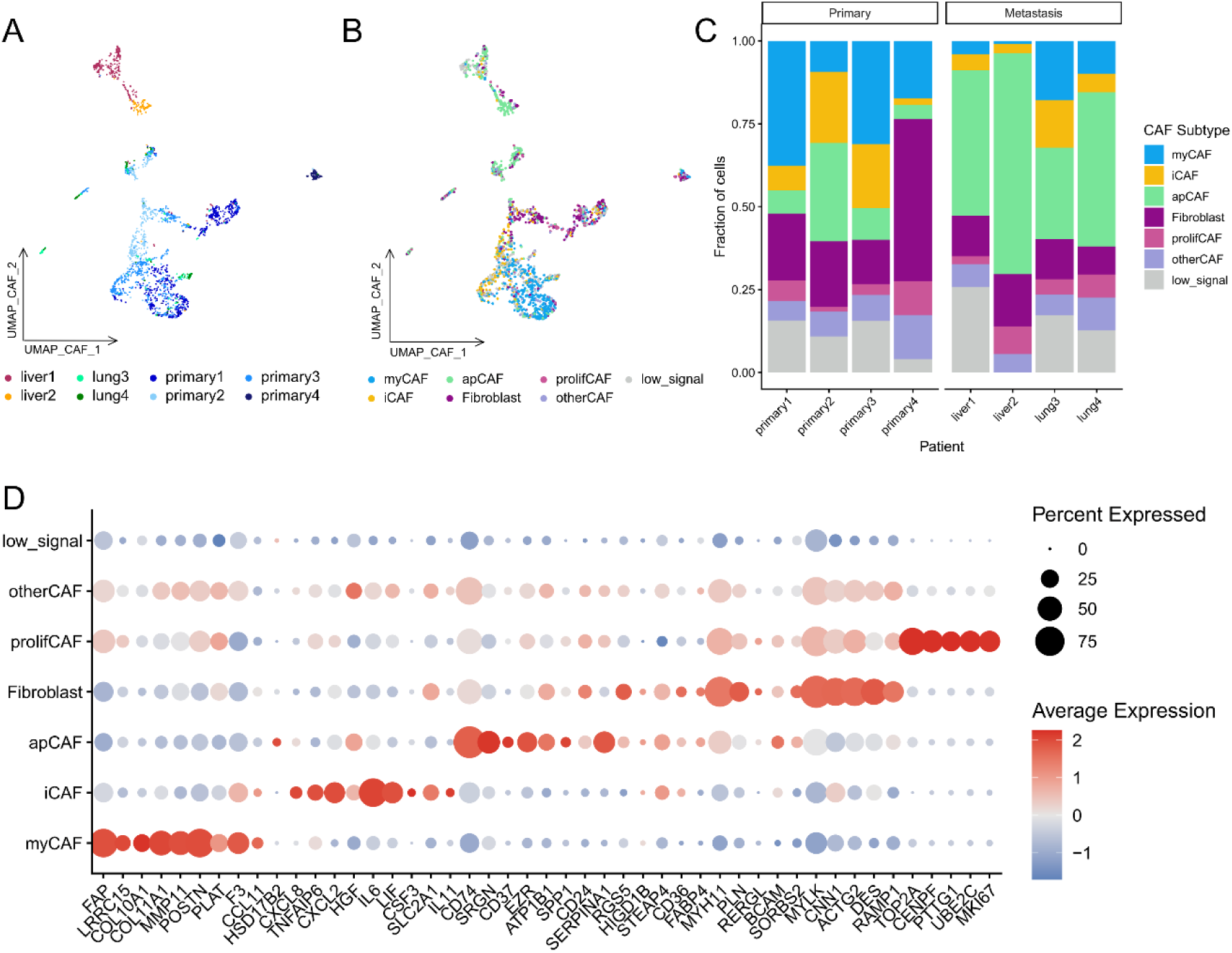
Single-nucleus CAF subtype analysis reveals enrichment of apCAFs in metastatic samples. (a) UMAP embedding of re-clustered fibroblast/CAF cells, colored by sample. (b) The same embedding colored by assigned CAF subtype (apCAF, antigen-presenting; iCAF, inflammatory; myCAF, myofibroblastic; prolifCAF, proliferating; Fibroblast; otherCAF, ambiguous assignment; low_signal, no signature above threshold). (c) CAF subtype composition per sample, split by primary tumors and metastases. (d) Expression of CAF subtype signature genes across assigned CAF subtypes. Dot size indicates the fraction of expressing cells, color indicates scaled mean expression.

Using a score-based classification (see Methods), CAFs were assigned to inflammatory CAFs (iCAF), myofibroblastic CAFs (myCAF), antigen-presenting CAFs (apCAF), proliferating CAFs (prolifCAF) and a fibroblast-like state (Fibroblast). Cells whose scores did not exceed the prespecified activation threshold were labeled low_signal, whereas cells with two similar highest scores were labeled otherCAF (**Fig. 2b,d; Fig. S2d**). In the CAF-restricted embedding, subtypes occupied largely distinct but partially overlapping regions, and the presence of both low_signal and otherCAF fractions is consistent with a continuum of CAF states rather than mutually exclusive identities.

We next compared CAF subtype frequencies between primary tumors and matched metastases (**Fig. 2c**; **Fig. S2b,c,e**). Fibroblasts/CAFs represented a mean of 6.1% of cells in primary tumors (range 3.0%-8.4%) and 3.7% in metastatic lesions (range 1.5%-9.7%) (**Table S6**). Within the Fibroblast/CAF population, apCAFs were enriched in metastatic lesions relative to primary tumors, with a numerically greater proportion in liver than lung metastases. apCAFs accounted for a mean of 12.6% of CAFs in primary tumors (range 4.1% - 29.7%), compared with 46.2% of CAFs in metastatic lesions (range 27.6% - 66.7%) (**Table S6**). Among metastatic sites, apCAFs represented a mean of 37.0% of CAFs in lung metastases and 55.3% of CAFs in liver metastases. Conversely, myCAFs tended to be more abundant in primary tumors, whereas the remaining subtypes showed only modest differences between sites.

These findings identify apCAFs as a potential metastasis-associated stromal population in LMS and suggest site-specific remodeling of the microenvironment during metastatic progression. Given the small size of the snRNA-seq cohort, we next sought validation in a larger cohort with preserved tissue context.

### Transfer of single-nucleus annotation into spatial context

To extend the single-nucleus findings into spatial tissue context, we performed imaging-based spatial transcriptomics (Xenium, 10x Genomics) on a tissue microarray (TMA) comprising 69 tumor hotspot cores from 21 LMS patients, including 14 matched primary-metastasis pairs (**Fig. S3a,b; Table S7**). Of note, the TMA also comprised the four pairs of the snRNA-seq experiment. One to three cores were analyzed per patient and lesion; the intended minimal number of two cores per condition could not be reached for every patient because individual cores were lost during sectioning. Profiling used a panel of 427 genes, combining a 377-gene tissue and cancer panel with 50 custom-designed sarcoma- and immune oncology-associated markers (**Table S8**). After quality control, 313,218 cells were retained across all cores, with a mean of 4,540 cells per core (range 340–13,427). Spatially resolved clusters were first defined *de novo* by Louvain clustering. Several clusters were shared across most patients, whereas others were restricted to individual metastatic sites or single patients (**Fig. 3a,b; Fig. S3c,d**), revealing the inter-patient heterogeneity of LMS and indicating both shared and patient- or site-associated cellular patterns in LMS.

**Figure 3.**
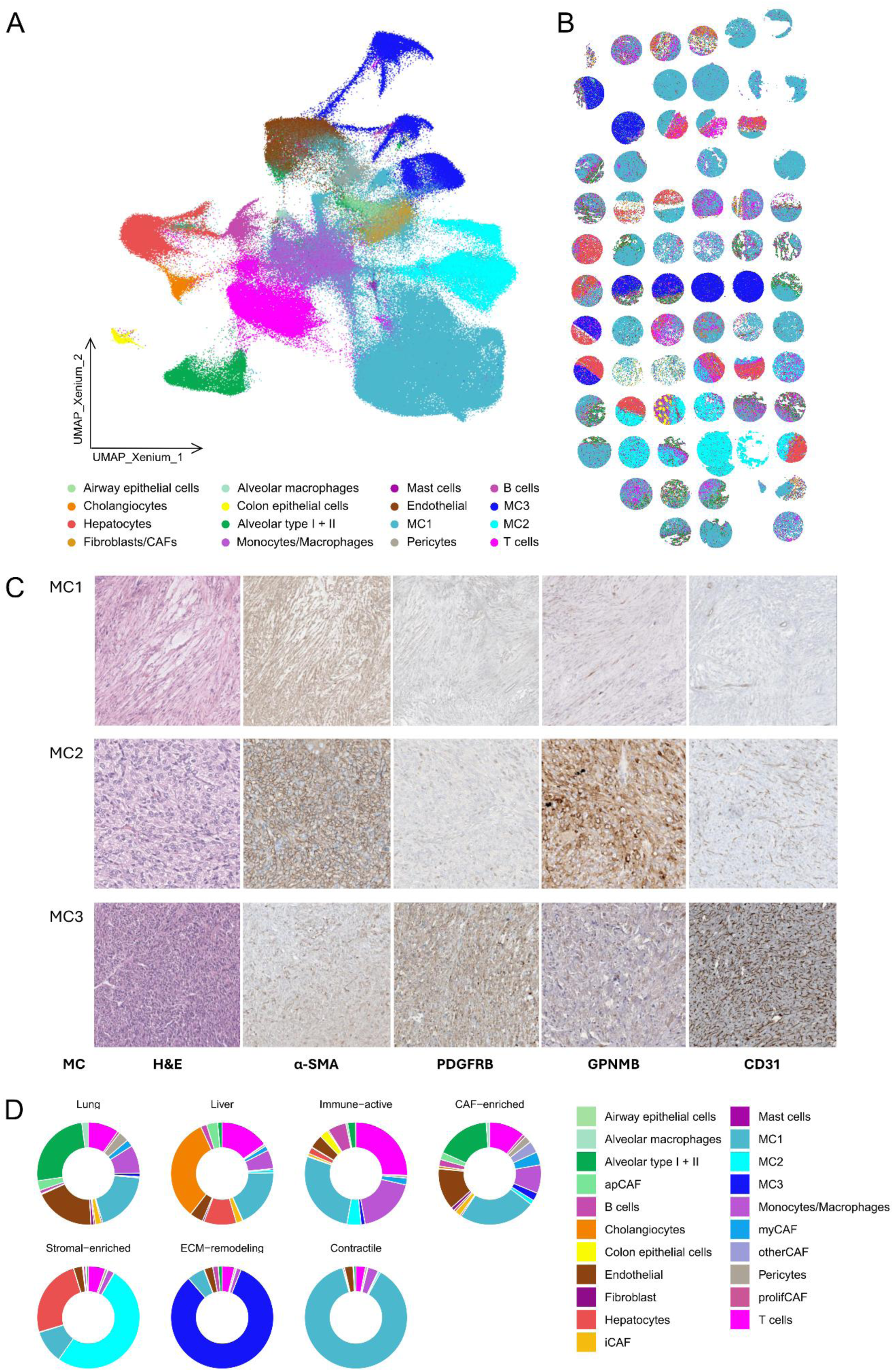
Spatial transcriptomics of the extended LMS cohort identifies three histomolecular phenotypes. (a) UMAP embedding of the spatial transcriptomics data annotated by transferred cell type labels (Seurat anchor transfer and SingleR, reviewed against HCE); MC1-MC3, malignant cell populations. (b) Spatial map of TMA cores colored by annotated cell type, colors correspond to those shown in (a). (c) Representative IHC staining for α-SMA, PDGFRB, GPNMB and CD31 for the three malignant clusters (MC1, MC2, MC3). (d) Cellular composition of the seven spatial niches identified by neighborhood analysis.

Transferring cell identities from the snRNA-seq reference proved challenging in this entity. Application of the widely used Robust Cell Type Decomposition (RCTD) algorithm left a large proportion of cells unannotated (∼63%; **Fig. S4a**), likely reflecting the limited transcriptional separation between malignant cells and stromal fibroblasts in this mesenchymal tumor together with the restricted gene panel in the spatial transcriptomics data that was designed towards epithelial malignancies. We therefore implemented a combined strategy integrating Seurat anchor-based label transfer with SingleR predictions (see Methods), and cell labels were critically reviewed by a pathologist (AM) in relation to the cellular cytology in coregistered Hematoxylin and eosin (HCE) stainings after Xenium imaging (**Fig. S5**). This approach annotated 14 cell types across the spatial transcriptomics cohort (**Fig. 3a,b**). Malignant cells were the dominant population, consistent with the tumor-focused selection of TMA cores, and stromal, immune and organ-resident populations of lung, liver and - in one core - colon were also resolved (**Fig. 3a,b**). Overall, malignant cells summarized per patient for localization accounted for an average of 62.4%, with proportions ranging from 21.2% to 97.1%. The mean number of malignant cells per patient was 5555, ranging from 118 to 18534 (**Table S9**).

### Spatial transcriptomics reveals three histomolecular leiomyosarcoma phenotypes

Within the extended spatial transcriptomics cohort (21 patients, 69 cores, **Table S7**) we next performed an intratumoral analysis. We could identify three major malignant populations termed malignant clusters 1-3 (MC1-MC3) (**Fig. 3a,b**; **Fig. S5**). MC1 was the predominant population and contained cells from the majority of patients (15 out of 21 patients, 49 out of 69 cores, **Fig. 3b**). In contrast, MC2 and MC3 each comprised cells from only three individual patients, indicating patient-restricted malignant states superimposed on a shared LMS tumor cell program (11 cores for MC2, 9 cores for MC3) (**Fig. S3d; Fig. 3a,b**). Marker expression distinguished the three populations: MYH11, consistent with a smooth muscle-like identity, was highest in MC1 and also present in MC2 but largely absent from MC3, whereas MC3 was characterized by FBN1 expression and MC2 by GPNMB (**Fig. S6a**). Outside MC2, GPNMB expression was generally low across other cell types but was also observed in alveolar macrophages (**Fig. S6b**). Comparative assessment of histomorphologies revealed that every malignant cluster harbored distinct morphological features (**Fig. 3c**): Within MC1 samples, myxoid growth features could be identified, MC2 displayed areas of high nuclear atypia with partially very large bizarre nuclei. HCE morphology of MC3 showed a dense capillary network, which was highlighted by CD31 immunohistochemistry (IHC) (**Fig. 3c; Fig. S7**).

We next assessed if the malignant cluster assignment could be captured by IHC staining. Candidates from routine pathology workup included α-SMA and PDGFRB, extended by GPNMB. IHC staining validated GPNMB-expression as overexpressed in MC2 while α-SMA and PDGFRB discriminated MC1 from MC3 (**Fig. 3c**). IHC stainings in whole-slide images confirmed a largely homogeneous protein expression across tumor tissue (**Fig. S8**).

Investigating cellular neighborhoods in the spatial transcriptomics data identified seven spatially distinct niches (**Fig. 3d**): two dominated by host organ tissue (lung and liver), a CAF-enriched stromal niche, an immune-active niche marked by T cells and monocytes/macrophages, and three malignant cell-dominated niches with stromal-enriched, ECM-remodeling or contractile composition. Notably, the three malignant-dominated niches contained different malignant populations - MC1 and MC2 predominated in the contractile and stromal-enriched niches, whereas MC3 was enriched in the ECM-remodeling niche (**Fig. 3c,d**). As for the association with clinicopathological parameters, we identified no association with prognosis (**Table S10**), although interpretation was limited by the small number of patients.

In summary, by multimodal integration of the snRNA-seq data and morphology with the spatial transcriptomics data, we identify the following histomorphologic phenotypes in LMS: (i) smooth-muscle-like with myxoid features, (ii) GPNMB⁺ with anaplastic features, and (iii) PDGFRB⁺ with vessel-rich stroma. Each phenotype could be identified with a concordant immunohistochemical (α-SMA, GPNMB, PDGFRB) and morphological correlate.

### Spatial validation of CAF subtype distribution and localization of apCAFs

For focused analysis of the CAF compartment, cells annotated as Fibroblasts/CAFs were selected for further analysis. Because the 427-gene panel lacked several of the marker genes required for score-based CAF subtype classification, CAF subtype labels were transferred from the snRNA-seq reference using SingleR. This recovered all CAF subtypes observed in the snRNA-seq cohort, including iCAFs, myCAFs, prolifCAFs, apCAFs, Fibroblast, otherCAF and low_signal cells (**Fig. 4a**). As in the snRNA-seq data, the transferred CAF labels occupied partially distinct but overlapping regions of the embedding, consistent with transitional CAF states.

**Figure 4.**
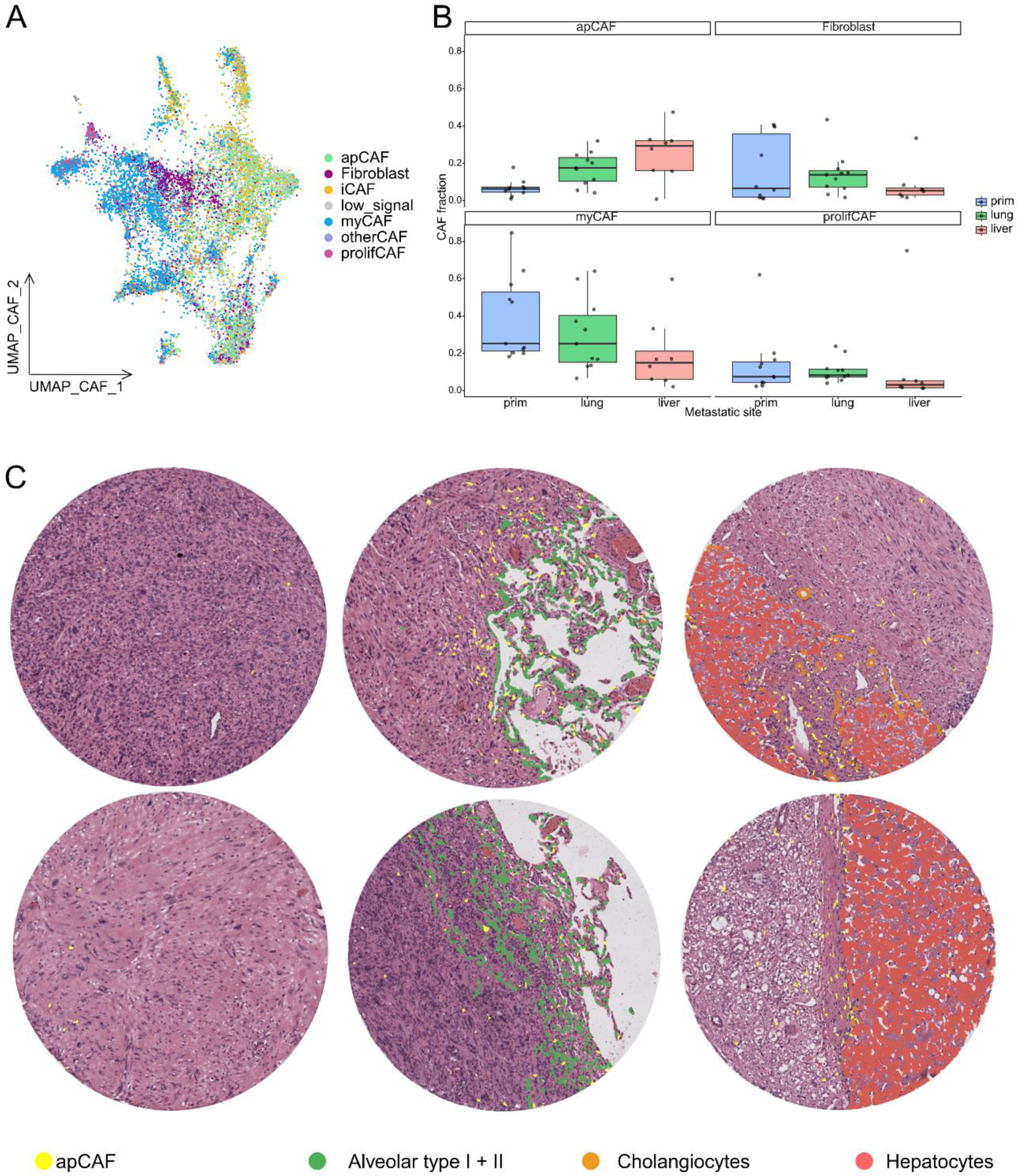
Spatial validation of apCAF enrichment and localization at the tumor–organ interface. (a) UMAP embedding of spatially profiled fibroblast/CAF cells colored by CAF subtype projected from the snRNA-seq reference by SingleR. (b) Fraction of apCAF, Fibroblast, myCAF and prolifCAF cells per patient, by tumor site (primary, lung metastasis, liver metastasis). Each point is one patient. (c) Representative HCE images with overlaid apCAF positions (yellow) in a primary tumor center, a lung metastasis and a liver metastasis; alveolar type I + II cells (green), cholangiocytes (orange) and hepatocytes (red) mark host organ tissue. Two examples per site.

The spatial cohort reproduced the compositional shift observed in the snRNA-seq data (**Fig. 4b**). At the patient-lesion level, apCAFs accounted for a mean of 6.1% of CAFs in primary tumors (range 0% - 17.8%), compared with 20.7% in metastatic lesions (range 0.82% - 47.4%) (**Table S11**). Effect sizes relative to primary tumors were large for both liver and lung metastases. The observed increase in apCAF frequency was associated with metastatic disease, as confirmed by an unpaired, two-sided Wilcoxon rank-sum test (p = 0.001). In patients with both a primary and a metastatic sample, the apCAF fraction was higher in the metastasis in 9 out of 10 cases (**Table S11**). Among metastatic sites, apCAFs represented a mean of 17.2% in lung metastases and 25.4% in liver metastases. Conversely, myCAFs tended to be more abundant in primary tumors (mean of 39.2%) than in metastases (mean of 25.4%). These compositional changes were concordant across the snRNA-seq and spatial cohorts. Per-patient compositions are shown in **Figure S9** and reveal individual fluctuations to the group-level pattern.

We next examined the spatial distribution of apCAFs in metastatic lesions. In both lung and liver metastases, apCAFs preferentially localize at the tumor–organ interface, forming a discontinuous band between malignant cells and the adjacent alveolar or hepatic parenchyma (**Fig. 4c**). In primary tumors, apCAFs were both sparse and showed no consistent spatial pattern. Interface localization was observed in cores from multiple patients and at both metastatic sites, supporting this pattern as a reproducible feature rather than a single-case observation.

Together, these findings demonstrate that apCAFs are enriched in the metastatic microenvironment of LMS and preferentially localize at the boundary between malignant cells and host organ tissue - the site at which tumor cells, host parenchyma and infiltrating immune cells are in closest proximity.

## Discussion

We present a paired single-nucleus and spatial transcriptomic characterization of primary and oligometastatic leiomyosarcoma. Profiling archival FFPE tissue made it possible to pair a primary tumor with a metastasis sometimes resected years later and combining dissociation-based with imaging-based single-cell transcriptomics allowed compositional observations from a small discovery cohort to be assessed in a larger cohort while preserving tissue architecture.

The principal finding is the enrichment of apCAFs in metastatic LMS, reproduced across two cohorts and two technologies, together with their localization at the tumor–organ interface. We use the term apCAF in its established sense as a transcriptionally defined CAF subtype; our data do not explicitly demonstrate antigen presentation. CAF subtypes remain largely unexplored in sarcoma, where the shared mesenchymal identity of tumor and stromal cells complicates the distinction between malignant and non-malignant fibroblasts^18^. The reciprocal enrichment of myCAFs in primary tumors and apCAFs in metastases indicates distinct stromal states across disease sites rather than a uniform expansion of fibroblasts: primary LMS may carry a more ECM-producing, myofibroblastic stroma, whereas metastatic lesions establish a spatially restricted apCAF-like niche.

apCAFs were first described in pancreatic ductal adenocarcinoma (PDAC)^16^, where lineage tracing showed that they can arise from mesothelial cells and induce naive CD4 T cells toward a regulatory rather than an effector phenotype^23^. A cross-tissue atlas subsequently resolved two apCAF populations present in most cancer types: mesothelial-like apCAFs located preferentially near cancer cells and fibrocyte-like apCAFs associated with lymphocyte-enriched niches^17^. The accumulation of apCAFs at the LMS tumor–organ interface matches the former pattern, although the number of apCAFs and the resolution of our data precluded robust separation of these states. Their position at the boundary where tumor cells meet host parenchyma and infiltrating immune cells suggests that they may help shape the local metastatic niche during colonization of a new organ, possibly by modulating immune interactions; this remains correlative, and the origin and function of apCAFs in LMS are open questions.

Assigning malignancy proved to be a recurring methodological obstacle that generalizes to sarcoma research. Reference-based annotation of the snRNA-seq dataset failed because malignant LMS cells and stromal fibroblasts share a mesenchymal transcriptional identity; in the spatial data the same problem manifested with the RCTD algorithm^24^ leaving approximately 63% of cells unannotated, consistent with benchmarks showing that annotation methods differ substantially on imaging-based platforms with small gene panels^25^. Multiple orthogonal solutions were required: inference of copy number profiles from expression data, established for separating malignant from non-malignant cells in solid tumors^26^ and here cross-validated against DNA methylation arrays for the single-nucleus dataset; and a combined anchor-based^27^ and SingleR transfer with pathologist review against coregistered HCE sections. Integrating single-nucleus and spatial transcriptomics with morphology was therefore essential for accurate cell-type annotation in this mesenchymal tumor.

The spatial data further resolved the malignant compartment into three histomolecular phenotypes, each defined by a transcriptional profile with a concordant immunohistochemical and morphological correlate. The predominant smooth-muscle-like phenotype with myxoid features (MC1) retains a differentiated smooth muscle program, with high MYH11 expression and strong α-SMA staining. The PDGFRB⁺ phenotype with vessel-rich stroma (MC3) shows the opposite pattern - low MYH11, absent α-SMA, strong PDGFRB and a dense CD31⁺ capillary network - indicating attenuated smooth muscle differentiation with acquisition of a perivascular or mesenchymal marker profile. Recurrent PDGFRB alterations have been reported in high-grade sarcomas with incomplete smooth muscle differentiation^28^; whether PDGFRB expression in this phenotype reflects genetic activation, a perivascular-like program or its ECM-remodeling niche was not assessed. The GPNMB⁺ phenotype with anaplastic features (MC2) occupies an intermediate position: α-SMA was retained in most cores and partially lost in a subset, consistent with attenuated but not lost smooth muscle differentiation, as described for poorly differentiated, pleomorphic LMS^29,30^. The three phenotypes thus likely span a gradient of smooth muscle differentiation, from retained (smooth-muscle-like, GPNMB⁺) to largely lost (PDGFRB⁺), and are separated immunohistochemically by GPNMB and PDGFRB, with α-SMA as a confirmatory marker. A recent spatial multi-omic study of LMS, available as a preprint, described a differentiated smooth-muscle-like and a dedifferentiated mesenchymal-like malignant state^31^; our smooth-muscle-like and PDGFRB⁺ phenotypes may correspond to these two ends of the spectrum, although a direct comparison would require the chromatin-accessibility data used to define those states.

The selective expression of GPNMB is of translational interest because GPNMB is a transmembrane glycoprotein with an antibody-accessible extracellular domain that has already been targeted clinically. GPNMB-directed antibody-drug conjugates (ADCs) and CAR T cells have therefore been explored as therapeutic strategies for GPNMB-expressing tumors. The GPNMB-directed ADC glembatumumab vedotin showed limited activity in recurrent osteosarcoma, indicating that target expression alone may not predict response^32^. GPNMB-directed CAR T cells showed potent activity against MiT/TFE-fusion-driven alveolar soft-part sarcoma and translocation renal cell carcinoma, with first-in-human evidence of tolerability and disease stabilization^33^, and a phase I trial in GPNMB-expressing solid tumors is recruiting (NCT07297667); an unbiased single-cell target discovery approach nominated GPNMB as a multi-cancer CAR T-cell target^34^; and in glioblastoma, GPNMB was identified as a dual-compartment antigen on tumor cells and immunosuppressive myeloid cells, with anti-GPNMB CAR T cells achieving durable disease control by depleting both^35^. GPNMB⁺ tumors were identified in three of 21 patients, which establishes the existence rather than the prevalence of this phenotype; its frequency in larger series, the homogeneity and membranous localization of expression, and its stability between primary and metastasis and under therapy remain to be established, for which GPNMB IHC as used here offers a straightforward assay.

Several limitations apply. The snRNA-seq cohort comprises four pairs, and comparisons within it are descriptive. The TMA consists of 1-mm cores, so cell type proportions are spatially sampled measurements rather than whole-tumor estimates. The GPNMB⁺ and PDGFRB⁺ phenotypes were each observed in three patients, and morphological features were assigned by pathologist review after cluster assignment; their recurrence and reproducibility require independent confirmation. While warranting further investigation, the presented cohorts do not allow further stratification by primary tumor site (e.g. uterine, retroperitoneal, extremities) due to cohort size and unbalanced groupings.

In summary, we provide a matched primary-metastasis reference for LMS and identify apCAFs as a metastasis-enriched stromal population at the tumor-organ interface and three histomolecular phenotypes recognizable by routine immunohistochemistry. Metastatic adaptation in LMS thus involves coordinated changes in malignant and stromal compartments shaped by organ context. Interface apCAFs and the GPNMB⁺ phenotype, the latter matching a surface antigen now entering CAR T cell trials, are candidate populations for functional and therapeutic investigation.

## Methods

### Study cohort

Eligible patients (age ≥ 18 years) had histologically proven metastatic leiomyosarcoma (LMS) and underwent surgical metastasectomy with curative intent. Clinical, histopathological, and outcome data were extracted from our prospectively maintained sarcoma database. Histopathologic diagnosis was reviewed by a specialized sarcoma pathologist. Synchronous metastasis was defined as metastatic disease occurring within six months of first diagnosis. Oligometastatic disease states were classified according to the ESTRO-EORTC classification by Guckenberger et al., allowing for a higher number of metastases if all lesions were considered amenable to local treatment^7^.

The study workflow is illustrated in **Figure 1a**. The cohort comprises 21 patients with LMS, of whom 14 are paired primary-metastatic cases. For generation of the snRNA-seq reference dataset, four pairs of primary tumors (3 retroperitoneal, 1 uterine) and matching metastatic lesions (two lung and two liver metastases) were selected. Selection criteria included high vital tumor content without major evidence of therapy-induced regression, sufficient tissue availability, and the presence of adjacent lung, liver, or stromal tissue.

The TMA-based spatial transcriptomics cohort includes all 21 patients (13 intraabdominal/retroperitoneal, 3 uterine, 5 extremities), with one to three cores per patient per condition. Clinical metadata, including date of initial diagnosis, time to metastasis, tumor grading, treatment and survival data were available for all patients. Survival analyses were performed from the date of first metastasectomy (**Table S3)**.

### Ethics

The internal review board and the ethics committee at the LMU University Hospital of Munich, Germany approved the study protocol (protocol no. 25-0373). In addition, the respective local ethics committees at each study site approved the current study.

### Single-nucleus RNA sequencing

Sample preparation for extracting the RNA was done in a tube-based format without xylene, with 2×25 µm thick sections for each tissue as described in the protocol (CG000784 Rev C). Viable cell count was determined with a Readycount Green/Red Cell Viability Stain (ThermoFisher, USA) on the Countess 3 Cell Counter (ThermoFisher, USA). Nuclei were extracted with an additional magnet bead-based purification step (Miltenyi, Ger), as described previously^21^. The further workflow including the determination of sample concentration and fragment length was performed with modifications to the hybridization process as described in Zhdanovich et al.^21^ and loading for the GEM chip according to protocol (CG000527, Rev F, Individual Wash Workflow). The samples were pooled after barcoding for the sequencing run.

Library concentrations were determined fluorometrically using a Qubit4 instrument (ThermoFisher) with the 1X dsDNA High Sensitivity Assay Kit (ThermoFisher, USA). Fragment length profiles were examined by capillary electrophoresis on an Agilent 1200 Bioanalyzer system using DNA High Sensitivity reagents (Agilent Technologies, USA). Libraries were adjusted to 2 nM in Illumina resuspension buffer. Sequencing was performed on a 300-cycle 10B flowcell on the NovaSeq X Plus Series (Illumina, USA) with a library loading concentration of 130 pM and 8% PhiX as control.

### DNA methylation array

The eight samples analyzed by snRNA-seq (P6414, P6427, P6501, P6470) were also subjected to DNA methylation analysis. For each sample, two 10 µm thick FFPE sections were used for nucleic acid extraction. Tumor areas were manually macrodissected from the sections using surgical blades based on HCE staining. Nucleic acid extraction was performed using the Maxwell RSC Blood DNA kit (Promega) on the Maxwell RSC 16 system (Promega, Germany) following an optimized protocol. DNA concentrations were quantified using a Qubit4 fluorometer (ThermoFisher).

Between 320 and 500 ng of genomic DNA per sample was subjected to bisulfite conversion using the EpiTect Fast Bisulfite conversion kit (Qiagen) according to the manufacturer’s instructions and as previously described^36^. DNA methylation profiling was conducted using the Infinium MethylationEPIC Bead-Chip array v2. Arrays were scanned using the NextSeq 550 platform (Illumina). Raw data processing and downstream analysis was performed as previously reported^37^.

### Generation of tissue microarray

On whole-slide HCE slides vital tumor areas were labeled by a pathologist (AM). With the TMA Grand Master (3D Histech, Hungary), depending on material availability, one to three punches with a 1 mm diameter were punched from each donor tissue block and transferred to the recipient block. To prevent individual cores from detaching and to ensure a uniform surface, the completed microarray was incubated at 42°C for 1 h on a glass slide and pressed flat before cutting.

### Imaging-based spatial transcriptomics

The generated TMA was sectioned at a thickness of 4.5 µm, hydrated in cold water followed by a 38°C water bath, and subsequently centered on a glass slide in accordance with Xenium, 10x Genomics platform requirements.

Slide processing was performed following the Xenium in situ workflow as specified in the manufacturer’s instructions (10x Genomics). A multi-cancer panel was applied together with a custom-designed 50-gene panel targeting sarcoma-associated genes and relevant immune checkpoint-related pathways (**Table S8**). Cell segmentation staining was performed prior to imaging on a 10x Genomics Xenium Analyzer (10x Genomics, USA; v4.0.1.4).

### Histochemistry and immunohistochemistry

HCE staining was carried out using an automated Tissue-Tek staining system (Sakura Finetek, Japan) in accordance with the manufacturer’s instructions.

For IHC, 2 µm sections from the TMA or FFPE whole-tissue blocks were deparaffinized and rehydrated using standard procedures. Heat-induced antigen retrieval was performed in citrate buffer (pH 6.0; 1x Target Retrieval Solution, Agilent Technologies, S2369) for 30 min in a microwave oven at 750 W. Endogenous peroxidase activity and non-specific binding were blocked using 7.5% hydrogen peroxide and 2.5% normal horse blocking serum, respectively (ImmPRESS anti-rabbit IgG Polymer Kit, Vector Laboratories, MP7401). Sections were incubated for 1 h at room temperature with an anti-GPNMB antibody (1:1500, clone E4D7P, Cell Signaling Technology). Antibody binding was detected using an anti-rabbit IgG polymer detection system (ImmPRESS, Vector Laboratories, MP7401) and visualized with DAB+ chromogen (Agilent Technologies, K3467). Slides were counterstained with hematoxylin, dehydrated, and mounted using Aquatex (Merck, 1.08562.0050). IHC staining for α-SMA (1:100, clone 1A4, Zytomed), PDGFRB (1:150, clone P-20, Santa Cruz) and CD31 (1:200, clone JC70A, Dako Agilent) was performed on the BenchMark XT (Ventana) automated slide stainer according to the manufacturer’s instructions.

### Analysis of snRNA-seq data

#### Preprocessing of snRNA-seq data

Demultiplexing was performed using the on-board BCLconvert software on the NovaSeq X Plus Series platform. Resulting FASTQ files were processed with CellRanger (10x Genomics, USA). Downstream analyses were conducted in R (version 4.4.2; released 2024-10-31) using Seurat (version 5.4.0, including SeuratObject version 5.3.0). Data were further processed in accordance with workflows from the Satija Lab^38^. Filtered feature-barcode matrices generated by Cell Ranger (version 9.0.1) were imported into R and processed using Seurat. Ambient RNA contamination was corrected using CellBender (Python 3.7.16, version 0.3.0), and the resulting denoised count matrices were used for downstream analysis. Cells were subjected to quality control filtering based on the number of detected genes and mitochondrial RNA content. Cells with fewer than 200 and more than 7,500 detected genes, or with mitochondrial gene content exceeding 5% were excluded from further analysis. Doublets were identified and removed using scDblFinder (version 1.20.2). Gene expression values were normalized using the SCTransform method. Dimensionality reduction was performed by principal component analysis (PCA) and uniform manifold approximation and projection (UMAP) embeddings were generated based on the first 30 principal components. Nearest neighbor graphs were constructed, and clustering was performed using the original Louvain algorithm with a resolution parameter of 0.15.

#### Cell typing

Cell clusters were identified using Louvain clustering in Seurat and initially annotated using multiple reference datasets. Annotation of lung cells was guided by a published lung cell atlas^39^ and the Azimuth lung reference^40^. Liver cells within liver metastasis were identified using the human liver cell atlas (https://www.livercellatlas.org/datasets_human.php). Initial tumor cell annotation was assessed using leiomyosarcoma and sarcoma reference datasets (GEO: GSM7634954; GEO: GSE212527^41^) (**Table S12**). However, malignant cell populations could not be reliably distinguished from reference datasets alone. Therefore, copy number variation (CNV) profiles were inferred by using inferCNV package (version 1.22.0), using monocytes/macrophages, endothelial cells, cytotoxic T cells, B and plasma cells and mast cells as non-malignant reference populations. Cells exhibiting pronounced CNVs were classified as malignant. Malignant classification was further supported by validation using DNA methylation data, where chromosomal gains and losses were consistent with inferCNV-based predictions. Final cell type annotations were validated based on canonical marker genes identified using the FindMarkers function in Seurat.

#### NMF analysis

Non-negative matrix factorization (NMF) was performed on malignant cells using the geneNMF framework (version 0.9.2) to identify transcriptional programs from SCT-normalized expression data. NMF was performed across factorization ranks k=4-9, with genes included based on a minimum expression threshold of 0.05. Individual gene programs identified across NMF runs were aggregated into metaprograms using cosine similarity-based clustering as implemented in the geneNMF framework, resulting in 20 initial metaprograms. Low-quality programs were removed based on manual inspection of silhouette score and number of genes, yielding a final set of 15 metaprograms for downstream analysis. Metaprogram activity scores were computed using the AddModuleScore_UCell function from the UCell package (version 2.10.1). Metaprograms were annotated using MSigDB Hallmark enrichment, manually curated gene signatures spanning major biological processes, and comparison with previously published Weizmann metaprograms^41^.

#### CAF subtyping

Non-negative matrix factorization (NMF) was performed on malignant cells using the geneNMF framework (version 0.9.2) to identify transcriptional programs from SCT-normalized expression data. NMF was performed across factorization ranks k=4-9, with genes included based on a minimum expression threshold of 0.05. Individual gene programs identified across NMF runs were aggregated into metaprograms using cosine similarity-based clustering as implemented in the geneNMF framework, resulting in 20 initial metaprograms. Low-quality programs were removed based on manual inspection of silhouette score and number of genes, yielding a final set of 15 metaprograms for downstream analysis. Metaprogram activity scores were computed using the AddModuleScore_UCell function from the UCell package (version 2.10.1). Metaprograms were annotated using MSigDB Hallmark enrichment, manually curated gene signatures spanning major biological processes, and comparison with previously published Weizmann metaprograms^42^.

### Analysis of spatial transcriptomics data

#### Preprocessing of spatial transcriptomics data

Xenium output images were imported into R using the Seurat LoadXenium() function. Quality control was performed by filtering transcript counts below 20 (nCount_Xenium) and above the 98^th^ percentile, based on inspection of the transcript count histogram. Gene expression matrices were normalized using Seurat’s default LogNormalize method. Dimensionality reduction was carried out using PCA and UMAP embeddings were computed from the first 30 principal components. Nearest neighbor graphs were constructed, and cell clustering was performed with the Louvain algorithm at a resolution of 0.17.

#### Label transfer and niche analysis

Cell type annotations were transferred from the single-nucleus reference dataset using a combination of anchor-based label transfer (Seurat) following the approach developed in the Satija lab, and SingleR package (version 2.8.0) from Bioconductor. The resulting annotations were refined based on marker gene expressions and manually confirmed by a pathologist (AM) using the HCE histology. The malignant cell compartment was subsequently subsetted from the spatial transcriptomics dataset and reclustered using the Louvain algorithm to resolve intratumoral heterogeneity. This analysis identified six malignant cell subclusters (MC1, MC2, MC3, MC4, MC5, MC6), which were further consolidated into three major malignant cell populations (MC1–MC3) based on marker gene expression profiles (**Fig. 3a,b; Fig S4b,c**). For characterization of malignant cell populations, 20 cluster-specific marker genes were identified using the Seurat FindMarkers function. For comparative differential expression analysis between the major malignant cell populations, count matrices were aggregated and analyzed using a pseudobulk approach. Differential expression testing was performed using DESeq2 by applying the DESeq workflow. Niche analysis was performed using the Seurat’s BuildNicheAssay() function, using neighbors.k = 30 and niches.k = 7 on the “fov” assay. CAF subtypes defined in the single-nucleus dataset were projected onto the spatial transcriptomics data using SingleR, and their spatial distribution was further assessed using Xenium Explorer (version 3).

### Statistics

Progression-free survival (PFS) and overall survival (OS) were analyzed using the Kaplan-Meier method and Cox proportional hazards regression. PFS was defined as the time from first metastasectomy to progression or death. OS was calculated from the first date of metastasis and the date of first metastasectomy to death. Patients without an event were censored at the date of last follow-up. Associations of clinicopathological variables with PFS and OS were assessed using univariable Cox proportional hazards regression and log-rank tests. Differences in apCAF proportions between primary tumors and metastases were assessed using a two-sided Wilcoxon rank-sum test. A p-value ≤0.05 was considered statistically significant. All statistical analyses were performed using R software (version 4.4.2; R Foundation for Statistical Computing, Vienna, Austria).

## Data and code availability

For the snRNA-seq data, the per-sample HDF5 expression matrices for all eight samples (‘.h5’) and the quality-controlled and annotated Seurat object (‘.rds’) including metadata have been deposited on Zenodo (https://doi.org/10.5281/zenodo.22254841) and are publicly available as of the date of publication. For the spatial transcriptomics data, the complete Xenium output directory containing the spatial transcriptomics data and associated files, the corresponding HCE stained TMA image (‘.ome.tif’) and the quality-controlled and annotated Seurat object (‘.rds’) have been deposited on Zenodo and will be made publicly available upon full publication. The code used for analysis and figure generation is available on GitHub (https://github.com/mocklab/lms-met).

## Acknowledgments

LMB is supported by the Else Kröner-Fresenius Foundation (EKFS) under the Immuno Oncology and Local Intervention (IOLIN) research program at the LMU University Hospital Munich. We thank Dr. Ina Dewenter for help in creating Fig. 1a. The research work of AM is supported by the Max-Eder Research Group Program of the German Cancer Aid (Deutsche Krebshilfe) and the Bavarian Cancer Research Center (BZKF).

## Author contributions

Conceptualization, LHL, ABM, AM; Methodology, AK, LB, AM; Software: AK, AM; Formal Analysis, AK, LB, JDH, ABM, AM; Investigation, AK, LB, ABM, AM; Resources, PJ, AM; Data Curation, AK, LB, AM; Writing – Original Draft Preparation, AK, AM; Writing – Review & Editing, AK, LB, TK, PJ, YZ, LMB, JDH, DK, RLZ, MH, DDG, MA, MG, MT, CM, FA, KF, PK, SF, FK, LHL, ABM, AM; Visualization, AK, LB; Supervision, AM; Project Administration, ABM, AM; Funding Acquisition, PJ, FK, ABM, AM.

