## Supplementary figures and tables for "Matched primary-metastasis single-nucleus and spatial profiling of leiomyosarcoma identifies antigen-presenting fibroblasts at the tumor-organ interface"

### Supplements

#### Supplementary figures

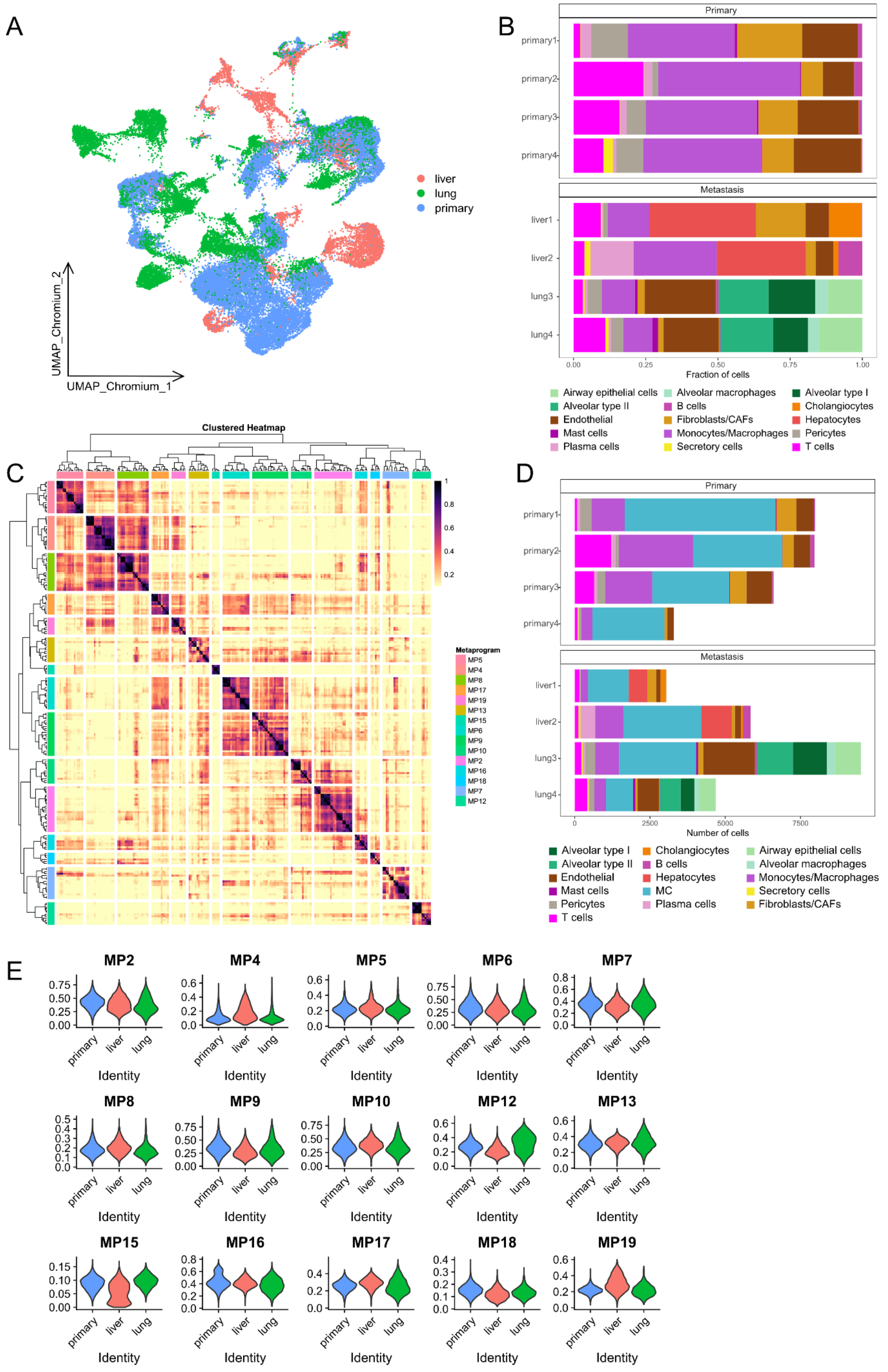

**Figure S1 - Single-nucleus RNA sequencing analysis of LMS samples.** (a) UMAP embedding of all single-nucleus samples showing shared and distinct cellular clusters across primary tumors, liver and lung metastases. (b) Relative amount of cell types across the eight single-nucleus samples without malignant cells. (c) Heatmap displaying the gene composition of identified metaprograms after quality control filtering. (d) Absolute cell numbers of identified cell types across the eight single-nucleus samples. (e) Metaprogram activity scores across malignant cells after quality control filtering.

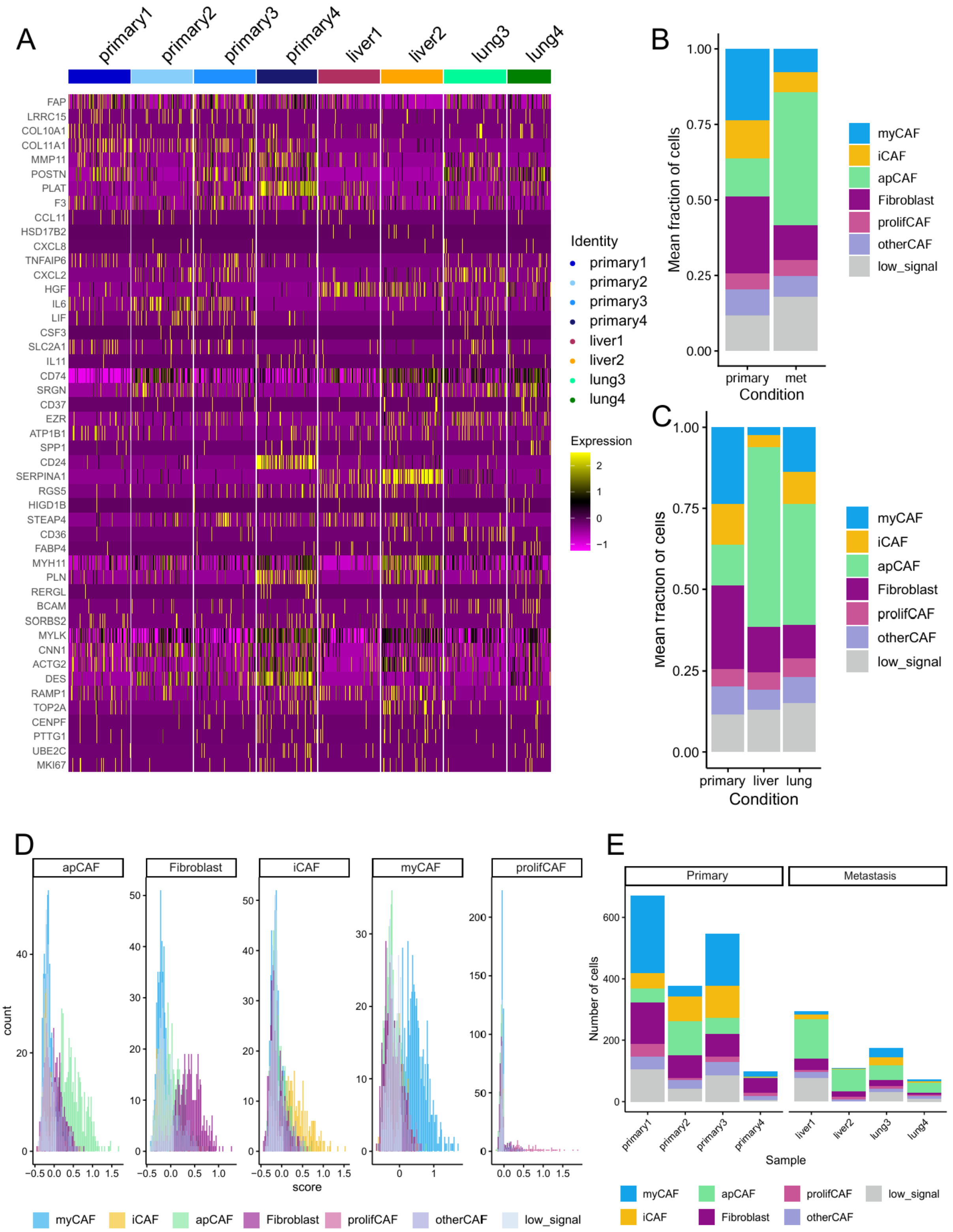

**Figure S2 - Characterization of CAF subtypes in the single-nucleus dataset.** (a) Heatmap showing expression of CAF subtype marker genes derived from Chen et al.[17]. (b, c) Mean fraction of CAF subtypes across primary tumors and metastatic samples, comparing (b) primary tumors and metastasis and (c) primary, liver and lung metastases. (d) Distribution of CAF subtype activity scores across all CAF cells, visualized as histograms for each identified CAF subtype. (e) Absolute numbers of cells assigned to each CAF subtype across the eight snRNA-seq samples.

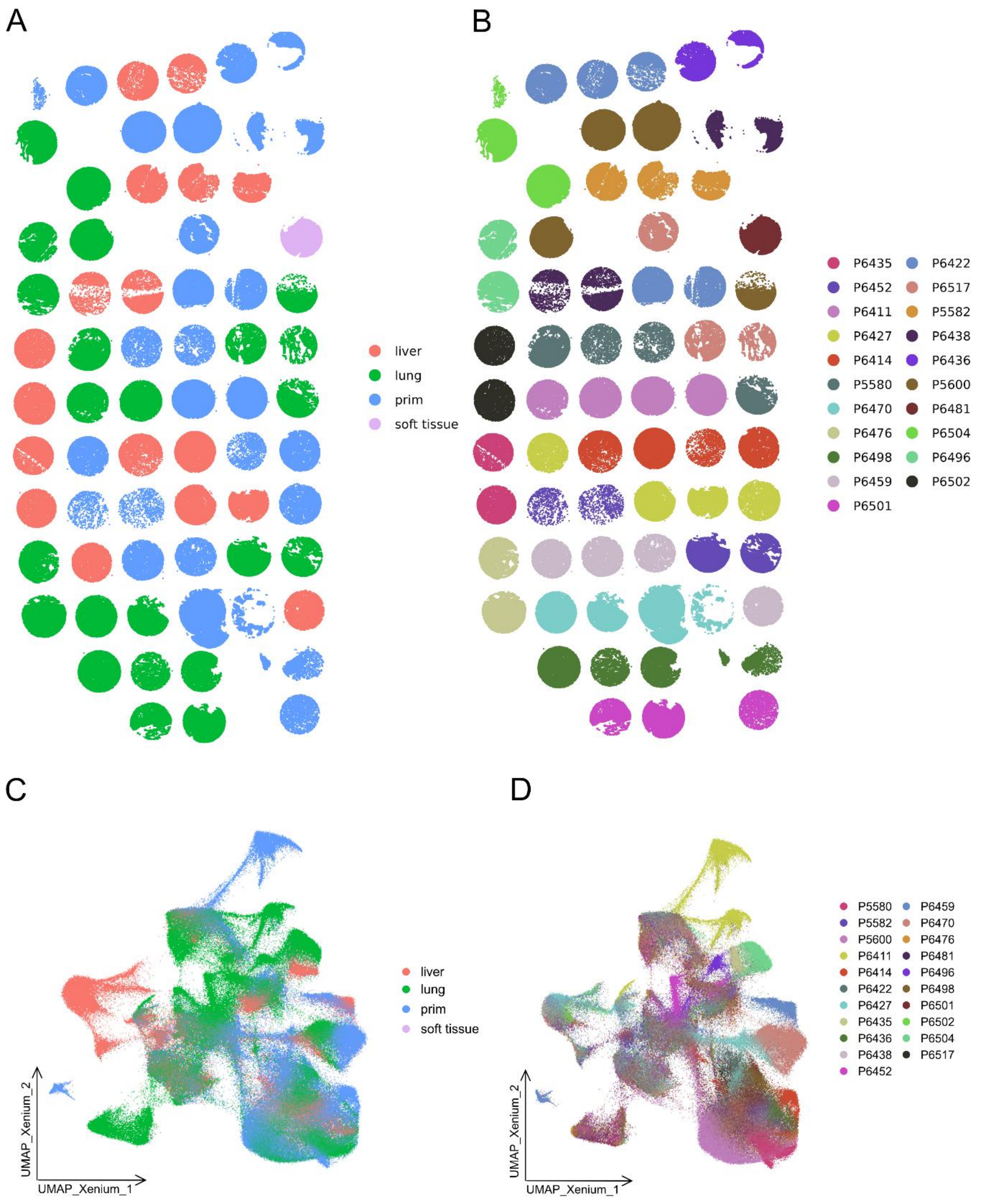


**Figure S3 - Spatial transcriptomics cohort overview.** (a) TMA core overview showing the anatomical distribution of primary tumors, liver, lung and soft tissue metastases. (b) TMA core overview displaying the distribution of patient samples within the spatial transcriptomics cohort. (c) UMAP embedding of spatial transcriptomic profiles, colored by anatomical site (primary tumors, liver, lung and soft tissue metastasis). (d) UMAP embedding of spatial transcriptomic profiles, colored by patient identity.

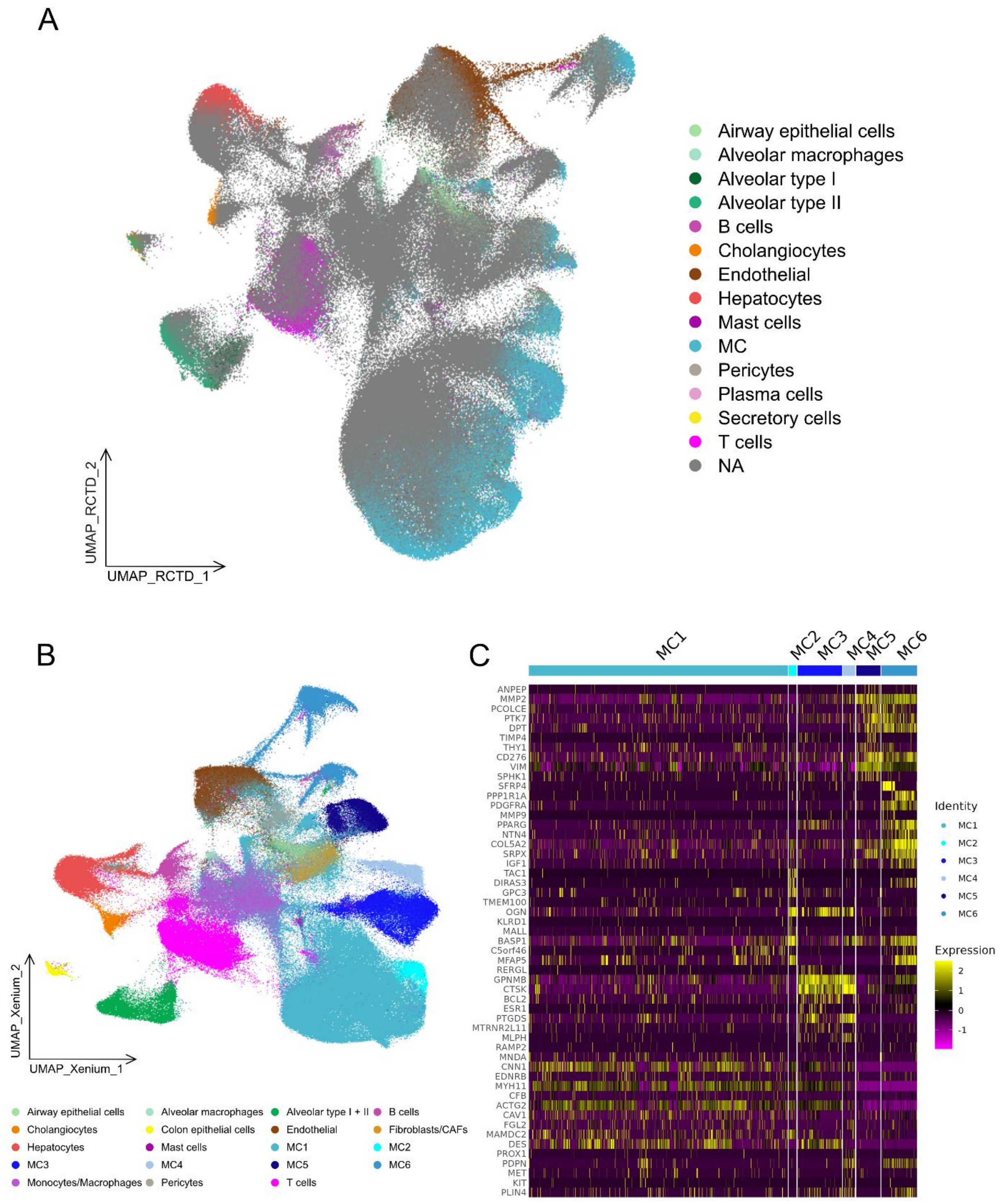


**Figure S4 - Label transfer and annotation of spatial transcriptomics data.** (a) RCTD-based label transfer resulted in a large proportion of cells remaining unannotated (NA). (b) Combined label transfer with SingleR and Seurat label transfer identified the major cell populations, including a malignant compartment that was further resolved into six distinct clusters by clustering-based annotation. (c) Expression of the top 10 marker genes across malignant clusters revealed distinct as well as partially shared marker expression patterns.

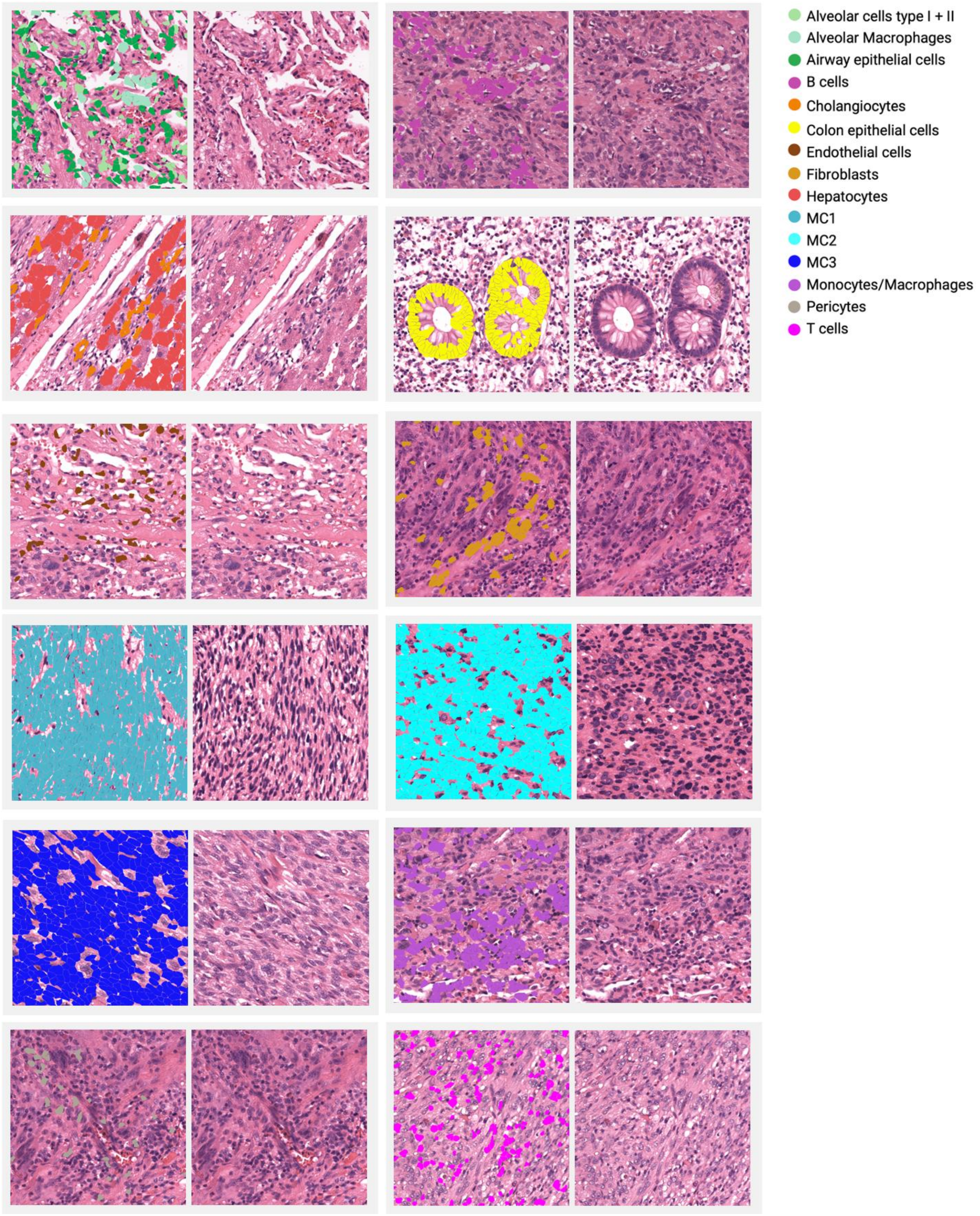


**Figure S5 - Final cell type annotation of spatial transcriptomics dataset.** Representative H&E images showing the final cell type annotation of the spatial transcriptomics dataset following label transfer and manual reannotation of selected clusters.

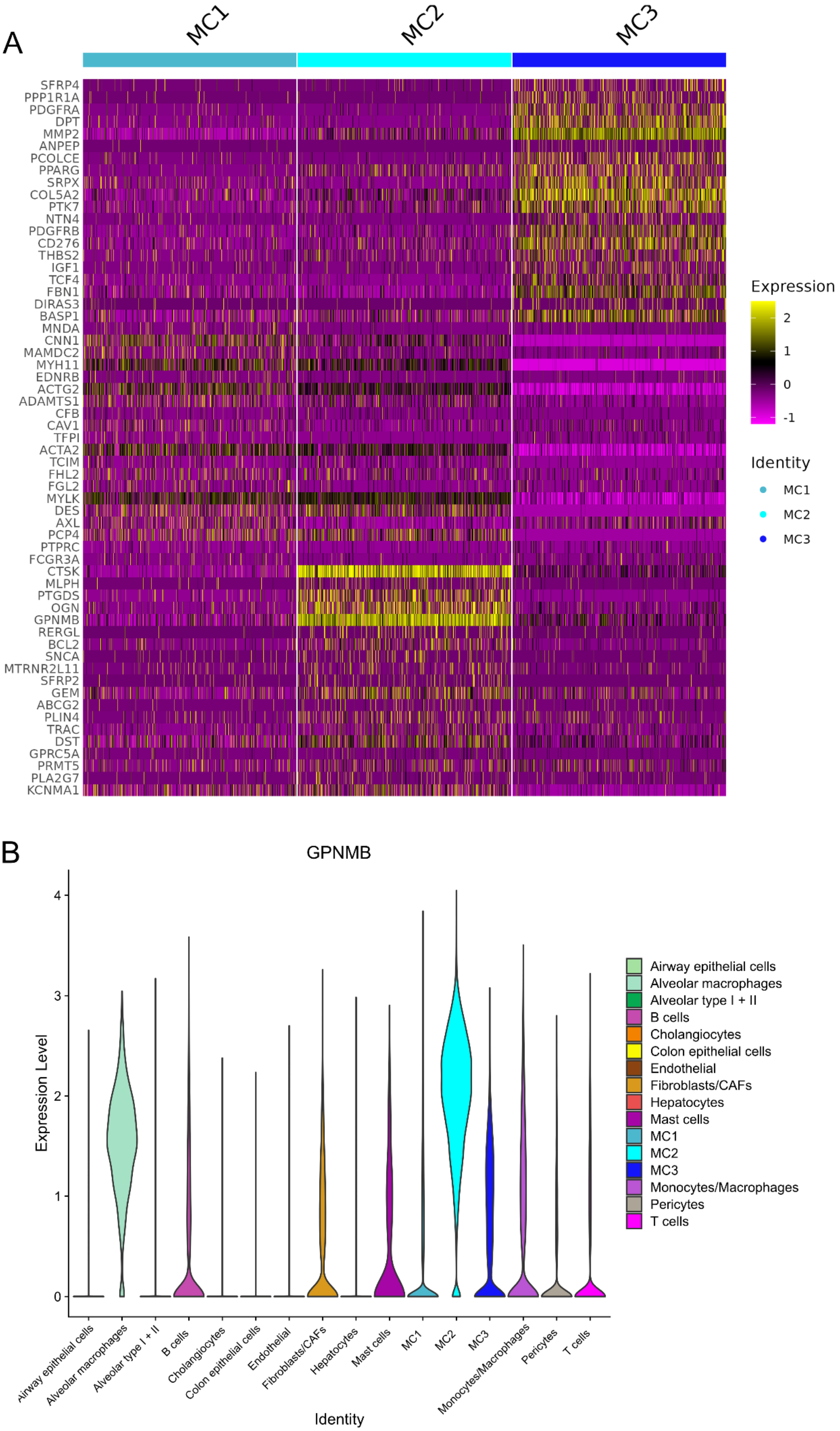
A
MC1
MC2
MC3
SFRP4
PPP1R1A
PDGFRA
DPT
MMP2
ANPEP
PCOLCE
PPARG
SRPX
COL5A2
PTK7
NTN4
PDGFRB
CD276
THBS2
IGF1
TCF4
FBN1
DIRAS3
BASP1
MNDA
CNN1
MAMDC2
MYH11
EDNRB
ACTG2
ADAMTS1
CFB
CAV1
TFPI
ACTA2
TCIM
FHL2
FGL2
MYLK
DES
AXL
PCP4
PTPRC
FCGR3A
CTSK
MLPH
PTGDS
OGN
GPNMB
RERGL
BCL2
SNCA
MTRNR2L11
SFRP2
GEM
ABCG2
PLIN4
TRAC
DST
GPRC5A
PRMT5
PLA2G7
KCNMA1
Expression
2
1
0
-1
Identity
MC1
MC2
MC3
B
GPNMB
Expression Level
4
3
2
1
0
Airway epithelial cells
Alveolar macrophages
Alveolar type I + II
B cells
Cholangiocytes
Colon epithelial cells
Endothelial
Fibroblasts/CAFs
Hepatocytes
Mast cells
MC1
MC2
MC3
Monocytes/Macrophages
Pericytes
T cells
Identity

**Figure S6 - Marker expression of the malignant clusters identified in the spatial transcriptomics dataset**. (a) Expression of the top 20 marker genes across the three identified malignant clusters in the spatial transcriptomics dataset. (b) Violin plot showing the distribution of GPNMB expression across the identified cell types, with pronounced expression in MC2. GPNMB expression was also observed in alveolar macrophages.

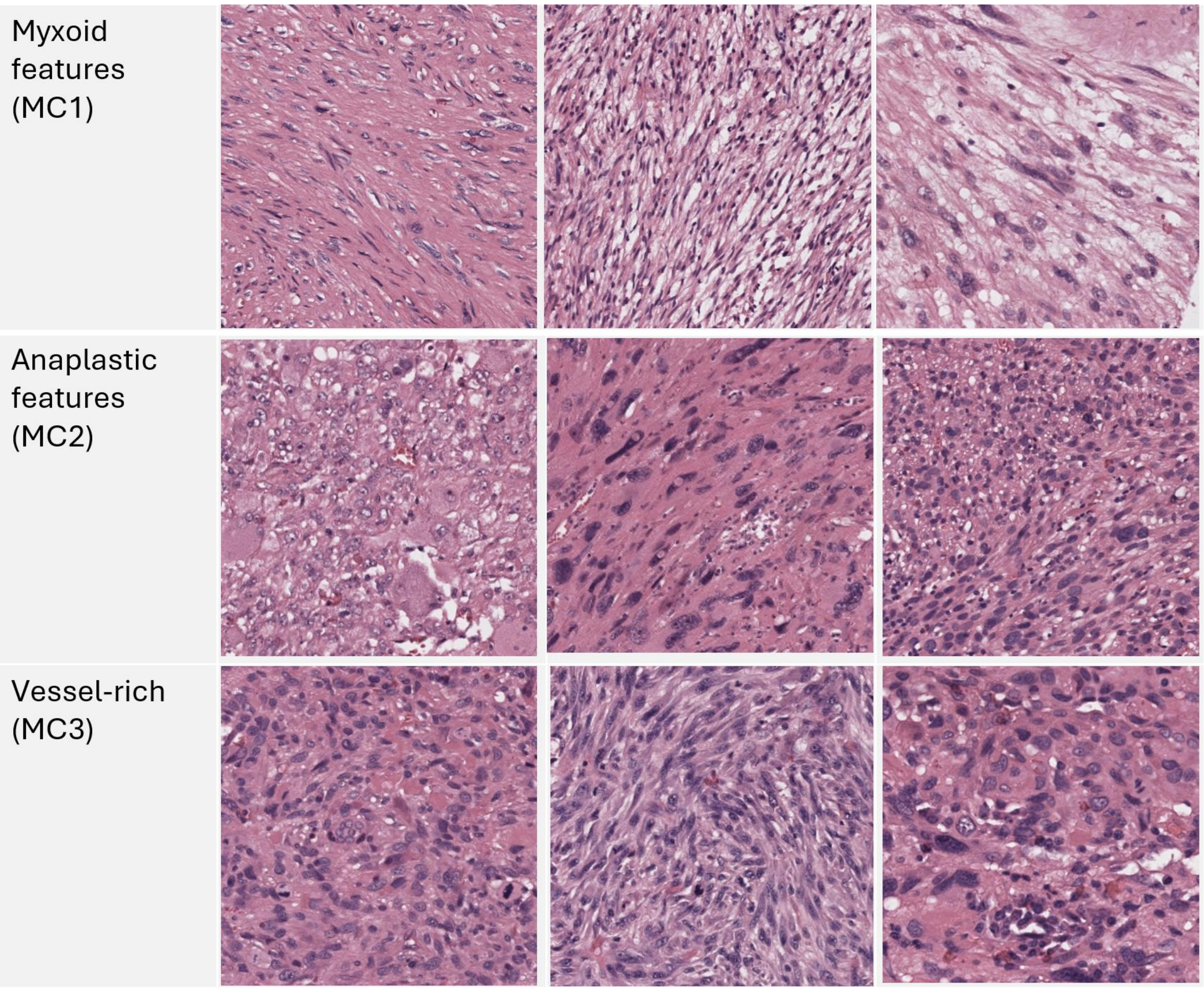


**Figure S7 - Distinct morphological features of the three malignant clusters.** (a) MC1 exhibits myxoid features, (b) MC2 shows anaplastic features, and (c) MC3 is characterized by a vessel-rich morphology.

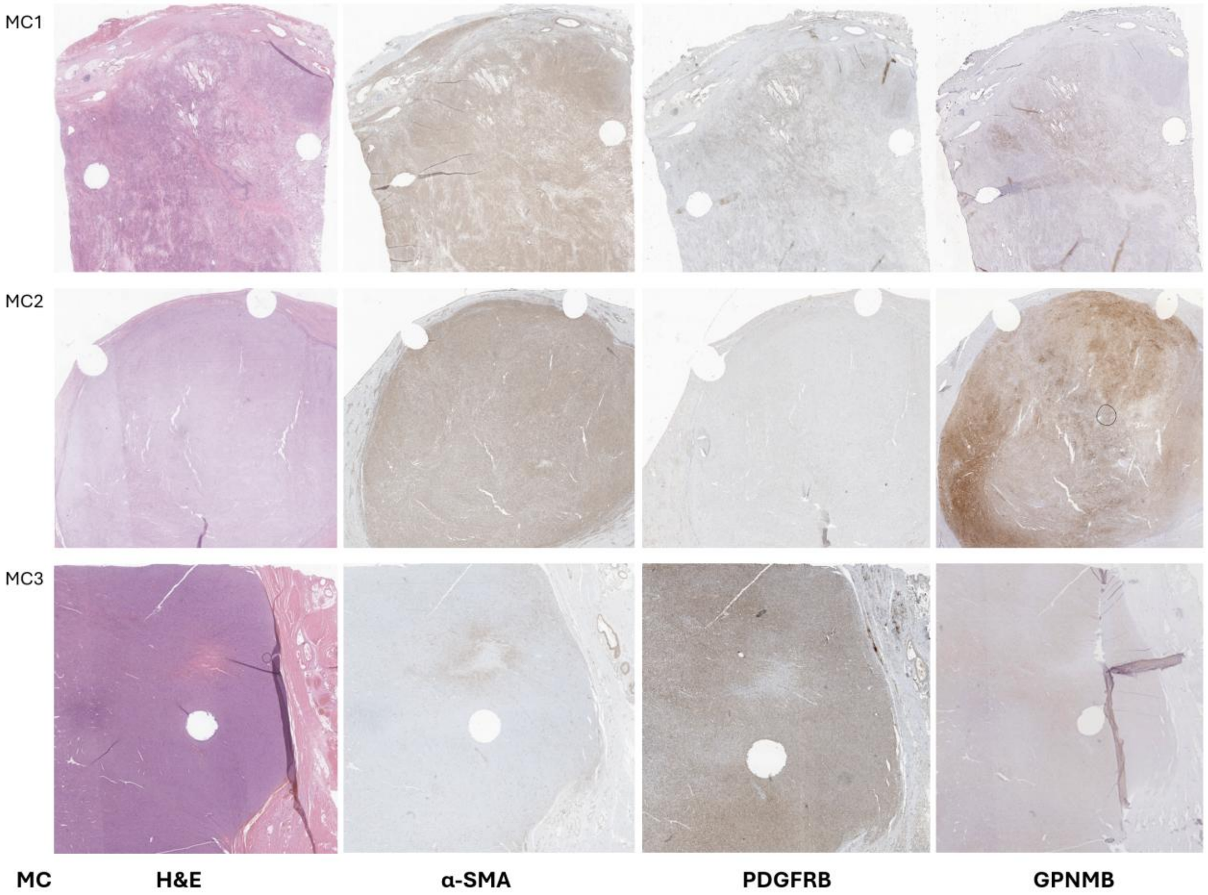


**Figure S8 - Whole-slide immunohistochemical validation of MC1-, MC2- and MC3-associated marker expression.** Representative regions for each malignant population (MC1, MC2, MC3). For each MC, corresponding H&E images and immunohistochemical staining for α-SMA, PDGFRB, and GPNMB are displayed.

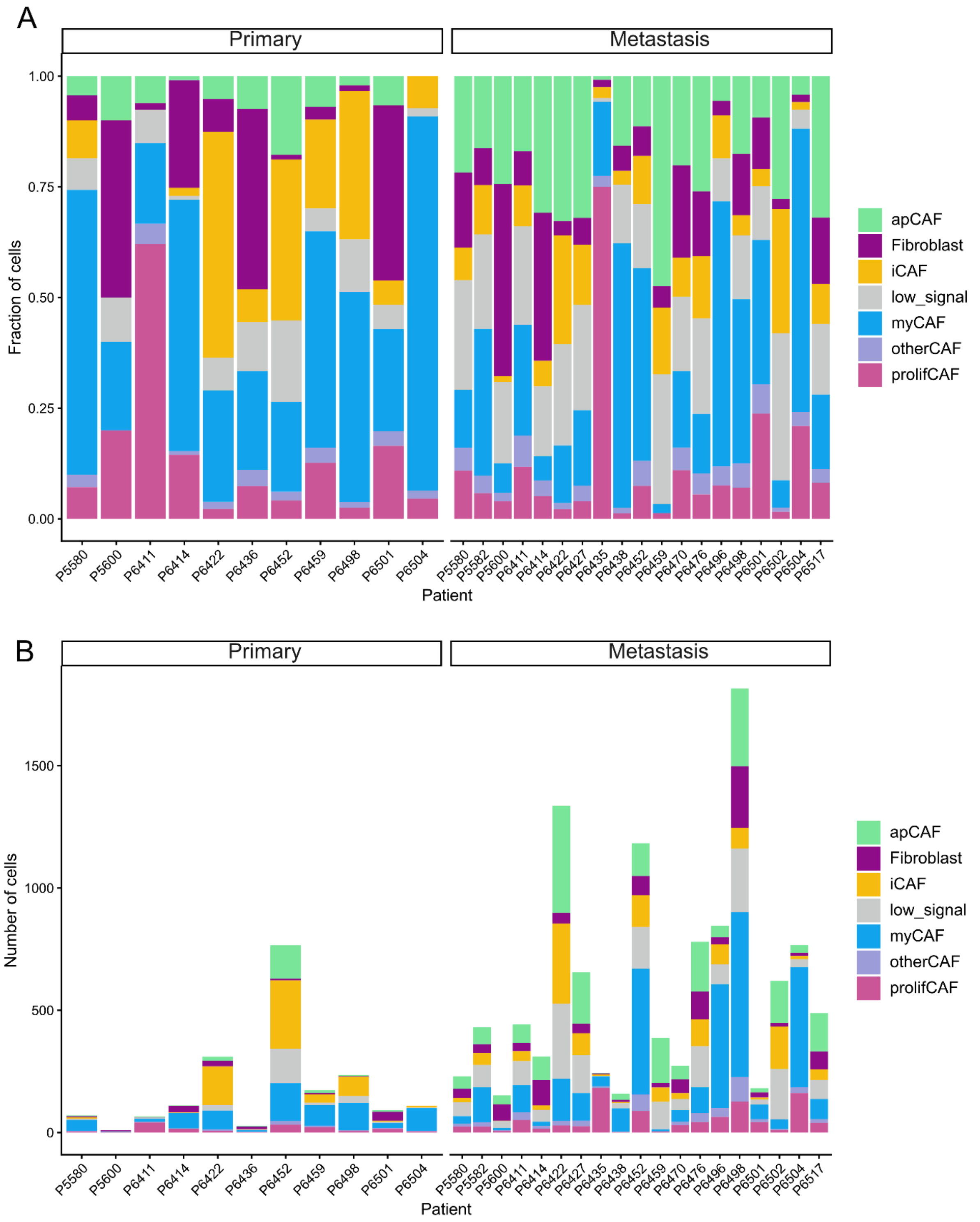


**Figure S9 - CAF distribution in the spatial transcriptomics dataset.** (a, b) (a) Relative proportions and (b) absolute cell numbers of CAF subtypes (apCAF, iCAF, myCAF, otherCAF, prolifCAF, Fibroblast, low_signal), stratified by tissue origin (primary versus metastasis) and patient.

### Supplementary tables

**Table S1 - Patient characteristics.** Clinical and treatment characteristics of the 21 patients included in the study, independent of biospecimen availability. Values are presented as number of patients (n) and percentage (%). NA, not available.

| **Factor** | **Value** | **n** | **%** |
|---|---|---|---|
| Total | | 21 | 100 |
| Age | Median: 54 years | | |
| Sex | Male | 4 | 19.0 |
| | Female | 17 | 81.0 |
| Grading | G2 | 13 | 61.9 |
| | G3 | 7 | 33.3 |
| | NA | 1 | 4.8 |
| Site of primary tumor | Extremities | 5 | 23.8 |
| | Intra-/retroperitoneal | 13 | 61.9 |
| | Uterus | 3 | 14.3 |
| Site of first metastasis | Lung | 12 | 57.1 |
| | Liver | 8 | 38.1 |
| | Intra-/retroperitoneal | 1 | 4.8 |
| Time to first metastasis | <12 months after first diagnosis | 10 | 47.6 |
| | ≥12 months after first diagnosis | 11 | 52.4 |
| First local therapy of metastasis | Surgical metastasectomy | 21 | 100 |
| Number of treated lesions | ≤5 | 17 | 81.0 |
| | >5 | 4 | 19 |
| Maximal diameter of treated lesions in cm | ≤2 | 10 | 47.6 |
| | >2 | 11 | 52.4 |
| Resection status | R0 | 18 | 85.7 |
| | R1 | 3 | 14.3 |
| Combination with systemic therapy | Yes | 11 | 52.4 |
| | No | 10 | 47.6 |
| Progression after first metastasectomy | Yes | 19 | 90.5 |
| | No documented progression | 2 | 9.5 |
| Deaths during follow-up | Yes | 7 | 33.3 |
| | No documented death | 14 | 66.7 |

**Table S2 - Available tumor specimens.** Clinical and pathological characteristics of the 35 tumor specimens included in the study for the 21 patients (see Table S1). Values are presented as number of specimens (n) and percentage (%). RHT, regional hyperthermia.

| Factor | Value | n | % |
|---|---|---|---|
| Total | | 35 | 100 |
| Tumor status | Primary tumor | 13 | 37.1 |
| | Primary tumor recurrence | 2 | 5.7 |
| | Distant metastasis | 20 | 57.1 |
| Specimen localization | Extremity | 4 | 11.4 |
| | Intra-/retroperitoneal | 11 | 31.4 |
| | Lung | 11 | 31.4 |
| | Liver | 8 | 22.9 |
| | Uterus | 1 | 2.9 |
| Pretreatment at time of specimen collection | None | 20 | 57.1 |
| | Chemotherapy +/- RHT | 12 | 34.3 |
| | Chemotherapy +/- RHT + radiotherapy | 3 | 8.6 |

**Table S3 - Prognostic clinical factors for progression-free survival and overall survival after first metastasectomy.** Univariable Cox hazards regression analysis to assess the association of clinical factors with PFS and OS. Hazard ratios are reported with 95% confidence intervals. For each variable, the first stratum listed was compared with the reference stratum. PFS, progression-free survival; OS, overall survival; HR, hazard ratio; CI, confidence interval.

| Variable | Strata | PFS HR (95% CI) | p | OS HR (95% CI) | p |
|---|---|---|---|---|---|
| Age at first diagnosis | >60 vs <=60 years | 0.31 (0.09-1.09) | 0.067 | 1.12 (0.22-5.79) | 0.893 |
| Sex | Female vs male | 1.16 (0.38-3.56) | 0.789 | 0.41 (0.08-2.15) | 0.291 |
| Histological grade | G3 vs G2 | 0.74 (0.27-2.00) | 0.548 | 1.19 (0.23-6.30) | 0.834 |
| Metastatic site | Lung vs other sites | 1.62 (0.63-4.14) | 0.317 | 0.94 (0.21-4.22) | 0.934 |
| Primary tumor site | Extremity vs other sites | 0.96 (0.34-2.72) | 0.942 | 0.74 (0.09-6.17) | 0.779 |
| Number of metastases | >5 vs <=5 | 2.47 (0.76-8.02) | 0.131 | 0.78 (0.09-6.49) | 0.816 |
| Largest metastasis diameter | >2 cm vs <=2 cm | 1.00 (0.39-2.56) | 0.998 | 2.30 (0.49-10.70) | 0.290 |
| Systemic therapy around metastasis-directed therapy | Yes vs no | 1.45 (0.54-3.89) | 0.459 | 3.71 (0.71-19.35) | 0.120 |
| Timing of metastatic presentation | Synchronous vs metachronous | 1.84 (0.69-4.90) | 0.220 | 3.68 (0.71-19.21) | 0.122 |

**Table S4 - Single-nucleus sequenced samples with counts and QC values.** snRNA-sequenced samples, including the total number of nuclei obtained, number of nuclei retained after quality control (QC), mean reads per nucleus, percentage of confidently mapped reads and year of FFPE block creation. Samples are identified by patient.

| Sample | Nuclei count before QC | Nuclei count after QC | Mean reads per nucleus | Confidently mapped reads | Year of FFPE block creation |
|---|---|---|---|---|---|
| primary1 | 8843 | 7986 | 9239 | 87.0% | 2017 |
| liver1 | 3395 | 3045 | 18198 | 72.4% | 2020 |
| primary2 | 8913 | 7966 | 6447 | 73.9% | 2016 |
| liver2 | 11967 | 5848 | 5744 | 78.6% | 2018 |
| primary3 | 7438 | 6610 | 10511 | 84.8% | 2019 |
| lung3 | 11217 | 9506 | 9757 | 68.4% | 2020 |
| primary4 | 3501 | 3294 | 27152 | 72.7% | 2015 |
| lung4 | 5180 | 4686 | 13506 | 84.6% | 2020 |

**Table S5 - Malignant cells in the snRNA-seq dataset.** Number and proportion of malignant cells identified in each snRNA-seq sample.

| Sample | Total cells after QC (n) | Malignant cells (n) | Malignant cells (%) |
|---|---|---|---|
| primary1 | 7986 | 5015 | 62.8 |
| primary2 | 7966 | 2963 | 37.2 |
| primary3 | 6610 | 2563 | 38.8 |
| primary4 | 3294 | 2395 | 72.7 |
| liver1 | 3045 | 1348 | 44.3 |
| liver2 | 5848 | 2593 | 44.3 |
| lung3 | 9506 | 2556 | 26.9 |
| lung4 | 4686 | 906 | 19.3 |

**Table S6 - Fibroblast/CAF abundance and apCAF frequency in the snRNA-seq dataset.** Number of Fibroblast/CAF cells and proportion of apCAFs within the Fibroblast/CAF population in each snRNA-sequenced sample.

| Sample | Total cells after QC (n) | Fibroblast/CAF cells (n) | Fibroblast/CAF cells (%) | apCAFs (% of Fibroblast/CAF) |
|---|---|---|---|---|
| primary1 | 7986 | 670 | 8.4 | 7.0 |
| primary2 | 7966 | 377 | 4.7 | 29.7 |
| primary3 | 6610 | 547 | 8.3 | 9.5 |
| primary4 | 3294 | 98 | 3.0 | 4.1 |
| liver1 | 3045 | 294 | 9.7 | 43.9 |
| liver2 | 5848 | 108 | 1.8 | 66.7 |
| lung3 | 9506 | 174 | 1.8 | 27.6 |
| lung4 | 4686 | 71 | 1.5 | 46.5 |

**Table S7 - Spatial arrangement of the TMA cores.** Each cell represents one TMA core (1 mm diameter) and indicates the tumor type and patient ID.

| | | | | | |
|---|---|---|---|---|---|
| prim, P6504 | prim, P6422 | prim, P6422 | prim, P6422 | prim, P6436 | prim, P6436 |
| lungmet, P6504 | | prim, P5600 | prim, P5600 | prim, P6438 | prim, P6438 |
| | lungmet, P6504 | livermet, P5582 | livermet, P5582 | livermet, P5582 | |
| lungmet, P6496 | lungmet, P5600 | | prim, P6517 | | softtissuemet, P6481 |
| lungmet, P6496 | livermet, P6438 | livermet, P6438 | prim, P6422 | prim, P6422 | lungmet, P5600 |
| livermet, P6502 | lungmet, P5580 | prim, P5580 | prim, P5580 | lungmet, P6517 | lungmet, P6517 |
| livermet, P6502 | lungmet, P6411 | lungmet, P6411 | prim, P6411 | prim, P6411 | lungmet, P5580 |
| livermet, P6435 | prim, P6427 | livermet, P6414 | livermet, P6414 | prim, P6414 | prim, P6414 |
| livermet, P6435 | prim, P6452 | prim, P6452 | livermet, P6427 | livermet, P6427 | prim, P6427 |
| lungmet, P6574 | livermet, P6459 | prim, P6459 | prim, P6459 | lungmet, P6452 | lungmet, P6452 |
| lungmet, P6476 | lungmet, P6470 | lungmet, P6470 | prim, P6470 | prim, P6470 | livermet, P6459 |
| | lungmet, P6498 | lungmet, P6498 | lungmet, P6498 | prim, P6498 | prim, P6498 |
| | | lungmet, P6501 | lungmet, P6501 | | prim, P6501 |

**Table S8 - Xenium custom-designed sarcoma- and immuno-oncology-associated markers.** List of custom-designed genes included in the Xenium assay in addition to the standard 377-gene expression panel.

| Gene | Gene | Gene | Gene |
|---|---|---|---|
| CTAG1B | AXL | STAT3 | MMP8 |
| CDK4 | LAMC2 | STAT6 | DDR2 |
| SOX10 | LAMB3 | TF | PRMT5 |
| MSLN | CALD1 | TLE1 | ITGB4 |
| CEACAM5 | INHBA | VIM | CSPG4 |
| ROR2 | MMP9 | CD276 | FGFR3 |
| NECTIN4 | DDR1 | ABCG2 | KDR |
| MAGEA1 | FGFR1 | BCL2 | PTK7 |
| MAGEA2 | GPNMB | BCL2L1 | SPHK1 |
| MAGEA3 | NFKB1 | TGFB1 | STAT1 |
| MAGEA4 | FN1 | JAK1 | STAT2 |
| TACSTD2 | MMP2 | JAK2 | |
| FGFR2 | PROM1 | JAK3 | |

**Table S9 - Malignant cells in the spatial transcriptomics dataset.** Number and proportion of malignant cells in the spatial transcriptomics dataset, stratified by patient and tumor localization.

| Patient | Tumor localization | Malignant cells (n) | Malignant cells (%) |
|---|---|---|---|
| P5580 | primary recurrence | 3365 | 78.29 |
| P5600 | primary | 18534 | 93.78 |
| P6411 | primary recurrence | 11842 | 95.29 |
| P6414 | primary | 5100 | 86.94 |
| P6422 | primary | 3719 | 39.09 |
| P6427 | primary | 5008 | 70.01 |
| P6436 | primary | 6458 | 90.55 |
| P6438 | primary | 4287 | 93.42 |
| P6452 | primary | 1026 | 54.09 |
| P6459 | primary | 4786 | 47.30 |
| P6470 | primary | 13821 | 94.03 |
| P6498 | primary | 1919 | 73.55 |
| P6501 | primary | 1806 | 70.66 |
| P6504 | primary | 118 | 34.71 |
| P6517 | primary | 2845 | 91.19 |
| P6422 | liver | 4352 | 50.40 |
| P5582 | liver | 6425 | 52.90 |
| P6414 | liver | 5601 | 54.42 |
| P6427 | liver | 3228 | 25.47 |
| P6435 | liver | 3379 | 51.70 |
| P6438 | liver | 2480 | 60.58 |
| P6459 | liver | 3602 | 42.58 |
| P6502 | liver | 1594 | 21.16 |
| P5580 | lung | 10664 | 77.64 |
| P5600 | lung | 8970 | 79.02 |
| P6411 | lung | 6551 | 56.91 |
| P6452 | lung | 4168 | 26.67 |
| P6470 | lung | 8047 | 64.32 |
| P6476 | lung | 6305 | 57.49 |
| P6496 | lung | 3737 | 37.09 |
| P6498 | lung | 5707 | 38.07 |
| P6501 | lung | 6824 | 66.36 |
| P6504 | lung | 10354 | 78.67 |
| P6517 | lung | 2145 | 31.75 |
| P6481 | soft tissue met | 5648 | 97.11 |

**Table S10 - Survival analysis according to malignant cell populations.** Association of malignant cell populations with PFS and OS following first metastasectomy. PFS, progression-free survival; OS, overall survival, MC, malignant cluster.

| Comparison | PFS HR (95% CI) | p | OS HR (95% CI) | p |
|---|---|---|---|---|
| MC2 vs MC1 | 0.72 (0.20–2.57) | 0.609 | 0.87 (0.10–7.27) | 0.900 |
| MC3 vs MC1 | 2.56 (0.64–10.17) | 0.181 | Not estimable | — |
| Global log-rank test (MC1 vs MC2 vs MC3) | — | 0.283 | — | 0.719 |

**Table S11 - apCAFs and myCAFs in the spatial transcriptomics dataset.** Number and proportion of apCAFs and myCAFs identified in each spatial transcriptomics sample, stratified by tumor type. Percentages represent the proportion of the respective CAF population among all CAFs identified in the corresponding sample.

| Patient | Tumor type | apCAF (n) | apCAF among CAFs (%) | myCAF (n) | myCAF among CAFs (%) |
|---|---|---|---|---|---|
| P5580 | Primary | 3 | 4.29 | 45 | 64.29 |
| P5600 | Primary | 1 | 10.00 | 2 | 20.00 |
| P6411 | Primary | 4 | 6.06 | 12 | 18.18 |
| P6414 | Primary | 1 | 0.90 | 63 | 56.76 |
| P6422 | Primary | 16 | 5.16 | 78 | 25.16 |
| P6436 | Primary | 2 | 7.41 | 6 | 22.22 |
| P6452 | Primary | 136 | 17.75 | 155 | 20.23 |
| P6459 | Primary | 12 | 6.90 | 85 | 48.85 |
| P6498 | Primary | 5 | 2.12 | 112 | 47.46 |
| P6501 | Primary | 6 | 6.59 | 21 | 23.08 |
| P6504 | Primary | 0 | 0 | 93 | 84.55 |
| P5580 | Metastasis | 50 | 21.74 | 30 | 13.04 |
| P5582 | Metastasis | 70 | 16.24 | 143 | 33.18 |
| P5600 | Metastasis | 37 | 24.34 | 10 | 6.58 |
| P6411 | Metastasis | 75 | 16.97 | 111 | 25.11 |
| P6414 | Metastasis | 96 | 30.87 | 17 | 5.47 |
| P6422 | Metastasis | 437 | 32.71 | 173 | 12.95 |
| P6427 | Metastasis | 210 | 32.01 | 112 | 17.07 |
| P6435 | Metastasis | 2 | 0.82 | 41 | 16.8 |
| P6438 | Metastasis | 25 | 15.72 | 95 | 59.75 |
| P6452 | Metastasis | 134 | 11.33 | 515 | 43.53 |
| P6459 | Metastasis | 183 | 47.41 | 8 | 2.07 |
| P6470 | Metastasis | 55 | 20.15 | 47 | 17.22 |
| P6476 | Metastasis | 203 | 26.03 | 105 | 13.46 |
| P6496 | Metastasis | 47 | 5.56 | 506 | 59.88 |
| P6498 | Metastasis | 318 | 17.51 | 674 | 37.11 |
| P6501 | Metastasis | 17 | 9.39 | 59 | 32.60 |
| P6502 | Metastasis | 172 | 27.74 | 38 | 6.13 |
| P6504 | Metastasis | 32 | 4.17 | 491 | 64.02 |
| P6517 | Metastasis | 156 | 31.97 | 82 | 16.80 |

**Table S12 - Reference datasets used for cell type annotation.** Summary of the publicly available snRNA-seq reference datasets used for cell type annotation, including the reference tumor type, data source, number of cells and number of annotated cell types.

| **Reference tumor type** | **Data source** | **Cell count** | **Number of cell types** |
|---|---|---|---|
| Liver | Liver Cell Atlas: Human | 167598 | 17 |
| Lung | Azimuth Human - Lung v2 (HLCA) | 584944 | 31 |
| Lung | Tabula Sapiens L | 15747 | 27 |
| Sarcoma | GEO GSE212527 | 3643 | 10 |
| Leiomyosarcoma | GEO GSM7634954 | 12736 | 3 |
